# Genomic and Metagenomic Analyses Reveal Parallel Ecological Divergence in *Heliosperma pusillum* (Caryophyllaceae)

**DOI:** 10.1101/044354

**Authors:** Emiliano Trucchi, Božo Frajman, Thomas H.A. Haverkamp, Peter Schönswetter, Ovidiu Paun

**Affiliations:** Department of Botany and Biodiversity Research, University of Vienna, Rennweg 14, 1030 Vienna, Austria; Institute of Botany, University of Innsbruck, Sternwartestraße 15, 6020 Innsbruck, Austria; Department of Biosciences, Centre for Ecological and Evolutionary Synthesis, University of Oslo, P.O. Box 1066 Blindern, 0316 Oslo, Norway

**Keywords:** Caryophyllaceae, E3 Ubiquitin Ligase, Microbiome, Phyllosphere, RAD sequencing

## Abstract

Cases of parallel ecological divergence in closely related taxa offer an invaluable material to study the processes of ecological speciation. Applying a combination of population genetic and metagenomic tools on a high-coverage RAD sequencing dataset, we test for parallel evolution across six population pairs of *Heliosperma pusillum* and *H. veselskyi* (Caryophyllaceae), two plant species found in the south-eastern Alps and characterized by clear morphological (glabrous vs. hairy) and ecological (alpine vs. montane, wet vs. dry) differentiation. Our analyses support a scenario of multiple independent instances of divergence between these species during the last 10,000 years. Structure analyses and simulations show that interspecific gene flow in each population pair is very low. A single locus, annotated as E3 ubiquitin ligase, an enzyme involved in plant innate immunity, shows a pattern of non-random segregation across populations of both species. A metagenomic analysis revealed information about contaminant exogenous DNA present in RAD sequencing libraries obtained from leaf material. Results of this analysis show clearly divergent bacterial and fungal phyllosphere communities between the species, but consistent communities across populations within each species. A similar set of biotic interactions is involved, together with abiotic factors, in shaping common selective regimes at different growing sites of each species. Different occurrences of *H. veselskyi* appear now genetically isolated from *H. pusillum* and from each other, and may independently proceed along the speciation continuum. Our work supports the hypothesis that repeated ecological divergence, observed here at an early stage, may be a common process of species diversification.

## Introduction

Repeated, independent instances of adaptive divergence, potentially leading to ecological speciation, constitute a rare opportunity to study the pace and mode of neutral and adaptive processes along the speciation continuum (Rundle & Nosil 2005; Schluter & Conte 2009; Roda *et al*. 2012). The early traces of selection on adaptive alleles are generally confounded by random drift that leads neutral or mildly deleterious alleles to fixation, especially in small populations. However, drift is expected to stochastically affect genomic loci along the entire genome, whereas divergent selection will produce hotspots of signals of reduced diversity and increased differentiation around only a few target regions (Cruickshank & Hahn 2014; Seehausen *et al*. 2014). Repeated evolution of ecotypes within the same taxon provides natural replicates that add significant power to tackle fundamental evolutionary questions about the trajectory of genomic divergence from locally adapted populations to isolated species, and to infer the adaptive role of different alleles, clearly disentangling the effects of neutral versus adaptive divergence (Lowry 2012; Seehausen *et al*. 2014).

Even if recurrent origin of morphological and/or ecological adaptation is a more common phenomenon than originally predicted (Levin 2001; Wood *et al*. 2005), only a few cases have been documented up to now. Besides the paradigmatic case of parallel freshwater adaptation in threespine stickleback (Colosimo *et al*. 2005), multiple origins of the same ecotype have been discovered among seaweeds (Pereyra *et al*. 2013), angiosperms (Brochmann *et al*. 2000; Berglund *et al*. 2003; Foster *et al*. 2007; Roda *et al*. 2013), invertebrates (Butlin *et al*. 2014; Soria-Carrasco *et al*. 2014), and vertebrates (Østbye *et al*. 2006). Parallel evolution can result from independent origins of the underlying molecular modification leading to a similar phenotype via a recurrent mutation at the same genomic location, through different alterations in the same gene producing a similar product, or through changes in different molecular components involved in the same phenotypic trait (see Stern 2013 for a review). Alternatively, and more commonly, similar selective pressures acting on different populations increase the frequency of adaptive alleles that are available as shared standing genetic variation, as a consequence of admixture among different populations or hybridization among species (Loh *et al*. 2013; Pearse *et al*. 2014). The result is collateral evolution *(sensu* Stern 2013) by shared ancestry as in the case of the repeated adaptation to freshwater habitats in the threespine stickleback (O’Brown *et al*. 2015), or collateral evolution by introgression as in the case of beak shape evolution in Darwin's finches (Lamichhaney *et al*. 2015).

In the early stages of ecological speciation, frequent gene flow may delay the appearance of reproductive barriers, and an emerging ecotype can ultimately be re-absorbed in the ancestral species (Levin 2004; Fant *et al*. 2010; Balao *et al*. 2015). Nevertheless, non-allopatric speciation with gene flow can produce genomic islands of strong differentiation scattered in a homogeneous and still largely recombining background (Feder *et al*. 2012; Martin & Orgogozo 2013). Divergent selection is then expected to maintain the adaptive features of the newly emerging ecotype even in the face of gene flow (Nosil *et al*. 2009). Other mechanisms grouped under the broad definition of “genomic conflicts” can, on the other hand, prevent admixture (Crespi & Nosil 2013; Seehausen *et al*. 2014). Processes like re-shuffling of genome architecture (Rieseberg 2001; Rebollo *et al*. 2010) or Bateson-Dobzhansky-Muller incompatibilities (BDMI; Orr 1995) have been suggested to strongly reduce gene flow during initial stages of speciation, favouring an early occurrence of reproductive barriers that can start as a reduction in hybrid fitness at one or both parental habitats (Presgraves 2010). In plants, for instance, a deleterious autoimmune response in hybrids has been related to the accumulation of BDMI (Bomblies & Weigel 2007; Chen *et al*. 2014; Sicard *et al*. 2015).

Ecological, morphological and genetic divergence in plants is also associated with consistent changes in the microbiome both at the level of the rhizosphere and, especially, the phyllosphere (Vorholt 2012; Bodenhausen *et al*. 2014). A different set of relationships with a novel range of microbial and fungal organisms can cause a feedback between the host plant and the associated microbial community (Bonfante & Anca, 2009; Lebeis 2014) fostering further divergence between populations thriving in contrasting habitats (Margulis & Fester 1991; Thomson 1999).

An excellent model to investigate repeated adaptive divergence is provided by the *Heliosperma pusillum* (Caryophyllaceae) species complex. This is a monophyletic group comprising perennial caespitose herbs that inhabit rocky habitats mostly on calcareous substrates in mountain ranges of southern Europe from the Sierra Cantabrica in the West to the Carpathians in the East (Frajman & Oxelman 2007). Members of the *H. pusillum* complex fall into two ecologically and morphologically distinct groups (Neumayer 1923; Frajman & Oxelman 2007; Frajman et al. 2009): i) a higher elevation (i.e., alpine) group occurring in damp, rocky habitats above the timberline, and ii) a lower elevation (i.e., montane) group inhabiting canyons and gorges as well as cave entrances and cliff overhangs with dry soils, but high atmospheric moisture and poor light conditions below the timberline. The higher elevation taxa are characterized by glabrous or sparsely hairy leaves and occasional presence of unicellular glands (Neumayer 1923; Frajman & Oxelman 2007), whereas plants of lower elevations share a dense indumentum with long multicellular sticky glandular trichomes. Dense indumentum has been explained as adaptation offering protection against drought, herbivores, and/or UV radiation (e.g., Levin 1973; Skaltsa et al. 1994; Espigares & Peco 1995; Kärkkäinen et al. 2004; Hanley et al. 2007) and genetic analyses indicate that the inheritance of this trait can be simple (e.g., in *Arabidopsis lyrata* a single recessive allele controls glabrousness; Kärkkäinen & Ågren 2002). In the *H. pusillum* complex, morphological variation is higher in the lower elevation group, which contains several narrowly distributed taxa (Frajman & Oxelman 2007; Niketic & Stevanovic 2007). Most of them are steno-endemics of the Balkan Peninsula, and only *H. veselskyi* extends its range to the south-eastern Alps. The few disjunct populations of *H. veselskyi* in this region occur nearby populations of the more widespread *H. pusillum* s. str. from the higher elevation group, even if the two species are always separated by several hundred meters of altitudinal difference, mostly inhabited by forests. Although the phenotypic differences between both taxa remain stable after cultivation under uniform conditions for at least three generations (C. Bertel & P. Schönswetter, personal observations), available phylogenies (Frajman & Oxelman 2007; Frajman et al. 2009) suggest that the relationships among populations of both species may be governed by geographic proximity rather than by ecological preference, implying that the disjunct populations of *H. veselskyi* potentially originated more than once from widespread *H. pusillum*.

Using genome-wide SNPs produced by restriction site associated DNA sequencing (RADseq), we investigate here the patterns of genomic divergence across multiple pairs of neighbouring populations of *H. pusillum* and *H. veselskyi* in the south-eastern Alps. We apply a combination of phylogenomic tools, analyses of population structure, outlier search and metagenomic approaches to test for parallel adaptation and understand the processes that shaped the recent evolutionary history of this group. We first ask if the observed genomic patterns support the hypothesis of multiple origins of the ecological divergence between higher vs. lower altitude species. We then investigate the extent and direction of gene flow between and within the two species. We also ask if the observed phenotypic divergence resulted from a similar genomic response to similar selective pressures by searching for adaptive loci, diverging in the same direction in different populations pairs. Finally, we ask whether the community of microorganisms in the phyllosphere of *Heliosperma* individuals is consistent across populations of the same species and if the marked divergence in ecology and morphology between the two species corresponds to different compositions of their phyllosphere.

## Methods

### Sampling and RADseq Library Preparation

Leaf material from ten individuals each was collected from six pairs of geographically close populations of *H. pusillum* and *H. veselskyi* (in the following termed “population pairs” and identified with uppercase letters from A to F) across the distribution range of *H. veselskyi* (Fig. 1, Table S1). Sampled material was dried and stored in silica gel. As the plants do not propagate vegetatively, the separation of individuals was straightforward.

**Figure 1.**
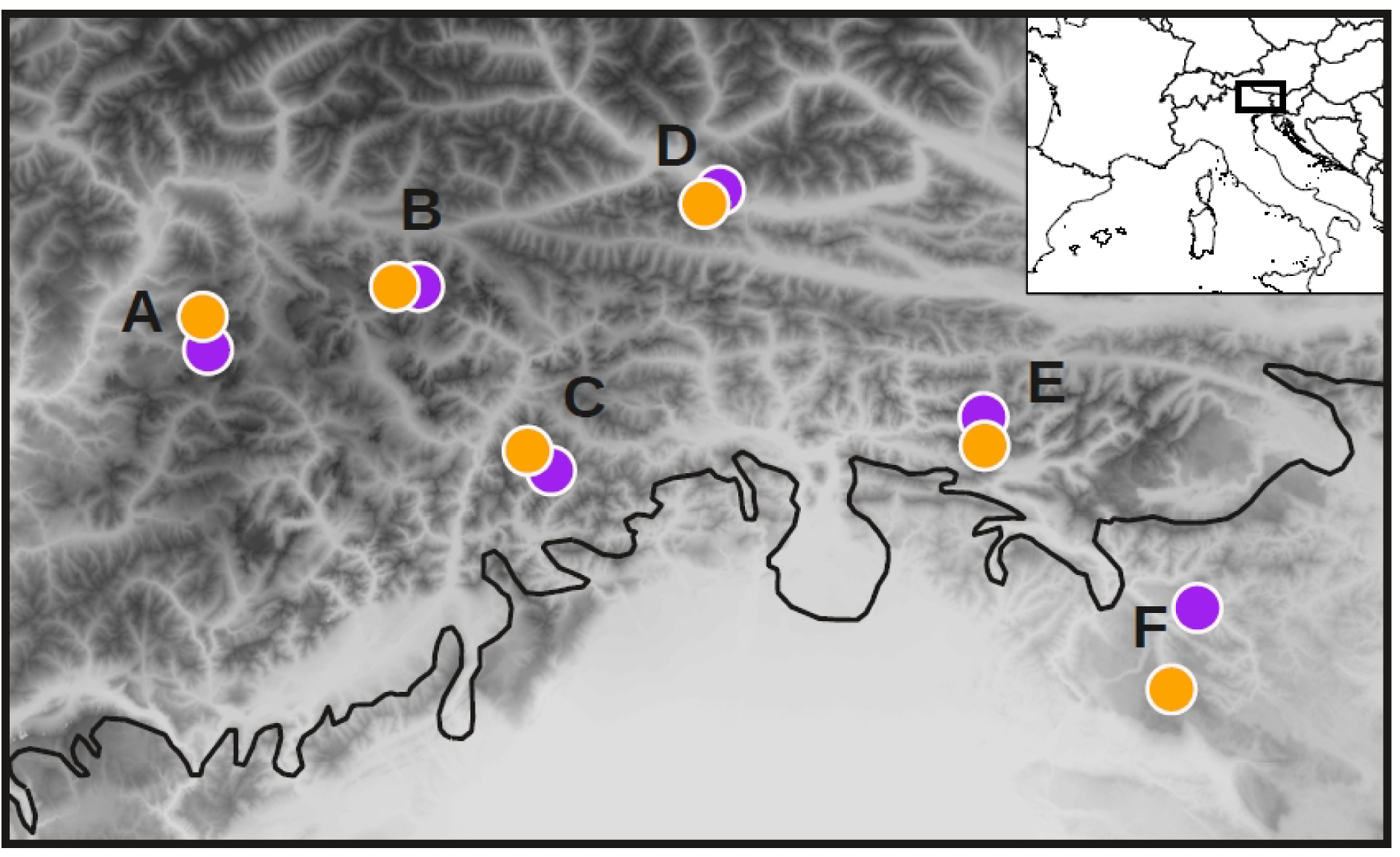
Map of sampled populations of *H. pusillum* (orange) and *H. veselskyi* (purple) in the south-eastern Alps. Extension of ice sheet during the Last Glacial Maximum is shown (black solid line). A: Grodental/Val Gardena (Dolomiten/Dolomiti, South Tyrol, Italy); B: Höhlensteintal (Dolomiten/Dolomiti, South Tyrol, Italy); D: Anetwände (Lienzer Dolomiten, Eastern Tyrol, Austria); C: Val Cimoliana (Alpi Carniche, Friuli, Italy); E: Sella Nevea (Alpi Giulie, Friuli, Italy); F: Idrijca valley (Primorska, Slovenia).

Single-digest RAD libraries were prepared using the SbfI restriction enzyme (New England Biolabs) and a protocol adapted from Paun *et al*. (2015) with modifications. In short, we started with 125 ng DNA per individual and ligated 125 mM P5 adapters to the restricted samples overnight at 16 °C. Shearing by sonication was performed with a Bioruptor Pico (Diagenode) with two cycles of 45s “on” and 60s “off” at 6 °C, targeting a size range of 200 to 800 bp. Four libraries were finally sequenced on an Illumina HiSeq at CSF Vienna (http://csf.ac.at/ngs/) as 100 bp paired-end reads.

### Identification of RAD Loci and SNP Calling

The raw reads were quality filtered and demultiplexed according to individual barcodes using the script *process_radtags.pl* included in the *Stacks* package (Catchen et al. 2013). As medium to large plant genomes largely consist of transposable elements (TEs), which may interfere with the locus-by-locus assembly in the *Stacks* analysis, we checked for the presence of TEs in the raw reads. For this aim, we blasted all paired-end reads, extending to the full library insert size, to the database of TEs identified in plants and collected in the RepBase database on the Giri repository (http://www.girinst.org/). Each individual sample was analyzed independently and the abundance of TEs (globally, and separately for the most common TE families) was compared among populations and between species (Supplementary Materials §2, Figs. S1–S2). However, as SbfI has a recognition site with 75% GC content, we expected a generally low representation of TEs, that are rather AT-rich (e.g., Turcotte et al. 2001).

The RAD loci were further assembled and SNPs were called using the *denovo_map.pl* pipeline in Stacks. A preliminary dataset (hereafter called *metagenomic dataset)* was built using a minimum coverage to identify a stack of 20* (-m 20), a maximum number of differences between two stacks in a locus in each sample of seven (-M 7), and a maximum number of differences among loci to be considered as orthologous across multiple samples of nine (-n 9). In this dataset, a bimodal distribution of the GC content across loci was discovered, with one peak below 45% GC and a second one above 60% (Fig. S3a). We suspected the latter to be due to bacterial DNA contamination, which is expected to result in a relatively high proportion of loci present in only a few individuals and having lower coverage than the loci of the target organism. We then increased our threshold for a locus to be assembled in each individual sample to 100 reads by setting the ‐m parameter in the *Stacks denovo_map.pl* pipeline to 100. Results then showed an unimodal distribution of GC content peaking at 40–45% (Fig. S3a).

The function *export_sql.pl* in the Stacks package was used to extract loci information from the catalog filtering for a maximum number of missing samples per locus of 60% and a maximum number of SNPs per locus of 10. Custom python scripts (Supplementary Materials §4) were used to further filter the loci for downstream analyses. A locus was discarded if i) the ratio of heterozygous individuals was > 0.65, an arbitrary cut-off used to reduce the risk of including paralogs in the dataset; ii) the GC content was > 65%, to further minimize the risk of including bacterial sequences; iii) any of the samples was scored as triallelic; iv) invariant. After checking the distribution of SNPs per position in the read, we observed an increase in the occurrence of SNPs in the last ten positions (Fig S3b). As we could not assess their validity, we decided to discard all SNPs found in this portion of the read. To check for the proportion of loci occurring on genes, filtered loci were blasted to NCBI nt database and to an available *Silene vulgaris* transcriptome (http://silenegenomics.biology.virginia.edu/: Sloan et al. 2011).

### Testing the Hypothesis of Parallel Evolution

First, in order to test for non-random accumulation of genetic divergence with increasing distance among populations, we tested for isolation-by-distance, both across all samples and each species separately, by comparing matrices of genetic and geographic distances. Nei's genetic distances among populations, including a sample size correction factor (Nei 1978), were estimated using the *gstudio* package in *R* (Dyer & Nason 2004). To assess the statistical correlation among matrices we applied Mantel tests with 9,999 randomizations.

We estimated a scenario of divergence among the investigated populations using the Bayesian coalescent approach implemented in *SNAPP* (Bryant et al. 2012). Our aim was to test if populations belonging to the same species were more closely related to each other than to the populations of the other species within the same population pairs, or vice versa. In the case of clustering by population pair, we were also interested in estimating the relative divergence time of each local split. As we were not interested in precise calibration of the tree but rather in understanding if the splits happened roughly at the same time in all population pairs, we applied a general mutation rate μ = 7 · 10^-9^ substitutions · site^-1^ · generation^-1^ (Ossowski et al. 2010) and explored the relative divergence time of each split. We randomly selected one biallelic SNP from each locus to create a data matrix where individuals were represented by one line each and each SNP position was coded as follows: 0, homozygote for the first allele; 1, heterozygote; 2 homozygote for the second allele. The model implemented in *SNAPP* assumes that all loci are unlinked, which is likely the case for most of our data (less than 1 marker/Mb considering a 2.2 Gb genome for this taxon). However, a moderate degree of linkage among markers is not expected to produce a strong bias in the results (Bryant et al. 2012). We set a prior for θ = 4μ*N*e as a gamma distribution with shape parameter (α) = 1.8 and an inverse scale parameter (β) = 5,000. Assuming a mutation rate μ = 7 · 10^-9^ substitutions · site^-1^ · generation^-1^ (Ossowski et al. 2010), our prior corresponds to a reasonable range of effective population sizes (95% HPD = 100-100,000), that should be large enough to take into account the small Ne for *H. veselskyi* populations and the generally larger Ne for *H. pusillum* populations (B. Frajman, field observation). We ran three independent MCMC chains in *SNAPP* for 2.5 · 10^5^ generations each, sampling every 100 generations. As SNAPP performance in terms of speed and convergence is dramatically reduced as long as sample size increases, we ran two additional analyses re-sampling three individuals at random for each population. In this case, the MCMC chains was run for 1 · 10^6^ generations. Convergence across different runs was checked in *Tracer v1.6* (http://tree.bio.ed.ac.uk/software/tracer/) and species trees were drawn in *DensiTree* (https://www.cs.auckland.ac.nz/~remco/DensiTree/).

The *fastStructure* algorithm (Raj et al. 2014) was used to estimate the population structure across all sampled individuals without *a priori* assumptions. This new algorithm, based on the same Bayesian model as the *Structure* software (Pritchard et al. 2000), implements a variational Bayesian framework and is particularly suited for analyzing large datasets of biallelic loci such as SNPs. It allows for testing an admixture model only. As only independent biallelic loci are allowed, we selected one SNP at random from each RAD locus. We first tested a flat beta prior distribution over population-specific allele frequencies at each locus (linear prior) using a range of *k* values (i.e., number of groups) from 2 to 12. The script *choose.py* included in the *fastStructure* package provides two estimates of the best *k:* one that maximizes the marginal likelihood, and a second estimate that best explains even very weak structure in the data. Both values for *k* were stored and the analysis repeated 100 times. Based on the results provided by the first round using the linear prior, we then refined our analysis using a logistic normal distribution with locus-specific mean and population-specific variance for the population-specific allele frequency (logistic prior in *fastStructure*) with values of *k* provided by the previous analysis (between 4 and 6 in our case). We replicated the analysis using the logistic prior 10 times and relevant values for *k* were recorded and summarized as before. The results of population structure for the best *k* were plotted using *python*.

The genetic similarity among all individuals was also investigated through a principal component analysis (PCA) using the *glPca* function in the *R* package *adegenet* (Jombart et al. 2011). The same analysis was replicated including only individuals from the westernmost localities A, B, C, and D that clustered tightly in the first analysis.

An analysis of pooled population variance was applied to measure the conditional graph distance (cGD) derived from population networks (Dyer et al. 2010). This algorithm is free of *a priori* assumptions about geographic arrangements of populations and uses a graph theoretical approach to determine the minimum set of edges (connections) that sufficiently explain the covariance structure among populations. Analyses and plotting were done in *R* using the packages *gstudio* and *popgraph* (Dyer & Nason 2004).

Using the *dist_amova* function in the *gstudio* R package and the *adonis* function in the *vegan* R package (Oksanen et al. 2013), a multilocus MANOVA was used to test for statistical differentiation among populations (i.e., 12 groups), population pairs (i.e., six groups) and species (i.e., two groups).

Global relationships among the individual samples were further investigated through a Maximum-Likelihood phylogenetic analysis. All loci were concatenated in a single sequence per sample coding heterozygosities as IUPAC ambiguities. The whole sequence in each locus was included in order to get empiric estimates of base composition and percentage of invariant sites. A Maximum-Likelihood algorithm with a GTRCAT substitution model was employed using 100 rapid bootstrap inferences in *RAXML 7.2.8* (Stamatakis 2006). Results were visualized and edited in *FigTree 1.4* (http://tree.bio.ed.ac.uk/software/figtree/).

### Assessing the Extent of Gene Flow between Species

To investigate the extent of local admixture between the two species, cluster analyses of the individuals collected from each population pair were performed using the *find.clusters* function in the *R* package *adegenet*. Similarly to *fastStructure*, this algorithm estimates the likelihood that the structure in the sample is described according to a range of clusters *(k-means* analysis ranging from 1 to 10 in our analysis) and then a Bayesian Information Criterion (BIC) is applied to choose the best *k*. The best *k* value was further tested using *fastStructure*. Separate analyses on population pairs were performed without prior information on taxonomic affiliation. The linear prior was applied and analyses replicated 20 times for each population pair. Results of population structure were plotted using *R* and *python*.

To investigate the rate of gene flow between the two species in each population pair we used a composite-likelihood maximization, by simulating joint-spectra under a continuous-time Markovian coalescent model in *fastsimcoal 2.5.11* (Excoffier et al. 2013). First, we estimated the folded 2-dimensions site frequency spectrum (2D-SFS) for each population pair. We then used the algorithm in *fastsimcoal2* that estimates parameters of interest in a demographic model through simulation of 2D-SFS and likelihood maximization. We built a simple demographic model with two populations that diverged at a certain time in the past and that continue to exchange individuals in both directions after the split. In this model, the two reciprocal migration rates were the only parameters to be estimated using a prior distribution from 0 to 0.1 (i.e., from 0 to 10% of the effective size of the receiving population). We then tested this model in each population pair. We fixed the divergence time for each population pair according to the SNAPP analysis and set a mutation rate μ = 7 · 10^-9^ substitutions · site^-1^ · generation^-1^. We set the extant effective population size to 5,000 for *H. pusillum* and 500 for *H. veselskyi* as a reasonable estimate of their difference in population size (B. Frajman, field observation). We ran 25 replicates of the same analysis for each population pair. For each replicate, we performed a maximum of 60 Expectation/Conditional Maximization (ECM) cycles over the two parameters (i.e., asymmetric migration rates between the two populations), each parameter optimisation step requiring the generation of 100,000 simulated joint-spectra. We then drew a distribution of the estimated parameters over the 25 replicates. Acknowledging that fixing the current population sizes was an arbitrary simplification, we tested the robustness of migration rate estimates with respect to different current effective population sizes. We then modified the model adding current population sizes as parameters to be estimated. We set a prior distribution between 100 and 30,000. Whereas the lower bound for the prior distribution is considered by the likelihood-maximization algorithms in *fastsimcoal2* as a strict limit, the upper bound is not strict. Although it resulted in unrealistically large values for the current effective population sizes, especially for *H. veselskyi* populations, the estimates of migration rates were always consistent with those obtained from the model with fixed population sizes.

### Detecting Outlier Loci

Using functions in the *R* package *adegenet*, we first explored the relative locus contribution to the principal component that explained the difference between species within population pairs. The *dapc* function was applied to provide an efficient description of the genetic clusters found by the *find.clusters* analysis. The contribution of each allele to the first principal component (always explaining the differentiation between species) was calculated applying the *loadingplot* function with a significance threshold of 0.01 (due to the higher global genetic divergence in F, a significance threshold of 0.005 was used in this population pair). The identity of alleles significantly contributing to the differentiation between species within population pairs was then compared among the six population pairs.

As the previous analysis was based on a selected dataset allowing only one random SNP per locus, we then tested the presence of outlier loci using the whole haplotype information with all SNPs phased for each locus. We used *Bayescan v.2.1* (Foll & Gaggiotti 2008) to search for loci with divergent allele frequencies between the two species. This analysis is based on F_ST_ and searches for outliers to the background of F_ST_ differentiation between the two groups of individuals compared. As we were interested in the adaptive differentiation shared by all populations of the same species and not in local differentiation within population pairs likely caused by random drift, we grouped all individuals according to species. *Bayescan* was run using a multistate character matrix with default options and results were plotted in *R*. The single emergent candidate locus was Sanger-sequenced in order to test its validity. A mini-contig was built using the paired-end reads of all individuals at that locus using *velvet* v.1.2.10 (Zerbino & Birney 2008). The mini-contig was blasted to the published *Silene vulgaris* transcriptome and annotated. Intronic and exonic regions were identified and several sets of primers were designed within and outside the coding region. We compared the results of amplification using only primers matching the non-coding part with amplifications obtained with primers matching only the coding region. A maximum-likelihood tree of all of the sequences amplified was built in *RAXML* using a GTRGAMMA model, and the tree was visualised with *FigTree*.

### Analysis of the Leaf Microbiome

We investigated the composition of the phyllosphere community employing the RADseq *metagenomic dataset* (see *“Identification of RAD loci and SNP calling”* above). The metagenomic analysis was performed on different sets of loci that were assembled in *Stacks* and selected in order to identify occasional (i.e., occurring in one individual) as well as more common (i.e., occurring in several individuals) taxa. In order to identify rare vs. common microbiome taxa, we performed the metagenomic analyses on three sets of loci in the *metagenomic dataset* selected to be present in at least one, five, or eight individuals, respectively, in one species and absent in the other. The analyses were implemented in two steps: *i*) the selected loci were blasted to the NCBI *nt* database using a maximum e-value of 0.0001 and *ii*) the positive hits were summarized using MEGAN (Huson et al. 2007). This software uses an algorithm to assign each hit to the lowest common ancestor (LCA) in the phylogenetic tree (preloaded from GenBank). Very conservative thresholds were applied in the LCA algorithm: to include a blast hit in the further analysis, a minimum alignment score of 100 (*min_score* = 100) was required, and to consider a taxon as identified, a minimum number of five hits (*min_support* = 5) had to be assigned to it, or to any of its descendants in the phylogenetic tree. Identified taxa were then summarized at the order level on the phylogenetic tree as identification at lower taxonomic level was considered less reliable. The datasets including loci present in at least five individuals of either species were further investigated recording number of individuals, coverage per individual, and number of populations in which each locus successfully assigned to a taxon by MEGAN was found. Finally, we also performed a metagenomic analysis on loci shared by the two species in each locality: the loci were selected when present in at least three individuals from both species. Besides confirming that most of the shared loci are actually assigned to the target *Heliosperma* species, this analysis allowed the identification of locally abundant taxa shared between the two species.

## Results

### Genomic Datasets

The total number of raw pairs of reads retained after quality filtering was ca. 255 millions with an average yield per sample of 2.1 (StDev = 0.8) million reads and an average PCR duplication rate of 2. The amount of paired-end reads mapping to the Viridiplantae TE database was in general very low for all individuals spanning from 0.46% (21,757 hits) to 4.1% (45,489 hits) with a slightly higher proportion (two samples Kolmogorov-Smirnov uncorrected *p*-value = 0.047) of hits to TEs found in *H. veselskyi* (Supplementary Information §2).

After *de novo* catalog building and SNP calling, we selected 1,719 loci present in at least 40 samples, with an average GC content between 40 and 45%. A total of 172 loci were discarded as invariant. The average coverage per allele was ca. 300* in *H. pusillum* individuals and ca. 200* in *H. veselskyi* (Fig. S3a). The lower coverage in *H. veselskyi* was due to higher exogenous DNA contamination in this species. It was possible to map 36% of the retained loci to the *Silene vulgaris* transcriptome and an additional 5% to coding sequences of other plant species in the NCBI database. Four of the loci matched two fungal sequences (poplar leaf fungus), one bacterial sequence *(Borrelia)* and one nematode sequence (Ascaridae). These loci were removed from the dataset. We further filtered our dataset according to the criteria detailed in the Methods section to a total of 1,097 variable loci, with an average of 102 individuals per locus, containing 3,401 SNPs (Fig. S3bc). This final dataset was used for all downstream analyses except the analysis of the phyllosphere communities.

### Testing the Hypothesis of Parallel Evolution

The isolation-by-distance model was confirmed for the global sampling (*p* = 0.001), as well as for each species analyzed separately (*H. pusillum*, *p* = 0.01, *H. veselskyi*, *p* = 0.02; Fig. 2). The Bayesian population tree reconstructed with *SNAPP* unambiguously showed at least five independent events of divergence between *H. pusillum* and *H. veselskyi* (Fig. 3) in the population pairs A, D, E, and F as well as in the unresolved B-C group. The time of divergence between the two species differs across localities: from the oldest split in F (roughly estimated as 10,000 years ago) to the most recent split in D (ca. 2,000 years ago). The *fastStructure* Bayesian analysis (Fig. 3) retrieved *k* = 5 as the optimal number of clusters. The composition of the five clusters closely matched the *SNAPP* population tree. Principal component analyses including all individuals, or only those from population pairs A, B, C, and D, clearly clustered individuals by population pair and not by species (Fig. S4). The groups and edges shown in the *popgraph* analysis (Fig. S5) generally corroborated the above results. In addition, this analysis suggested a tight connection among the three *H. pusillum* populations from B, C and D. The MANOVA showed that most variance is explained by populations and population pairs, but not by species (differentiation among 12 populations: R^2^ = 0.91, *p* = 0.001; six population pairs: R^2^ = 0.83, *p* = 0.001; two species: R^2^ = 0.02, *p* = 0.14). Finally, the maximum-likelihood tree reconstructed from the concatenation of all loci fully supported a scenario of parallel independent divergence between *H. pusillum* and *H. veselskyi* (Fig. S6).

**Figure 2.**
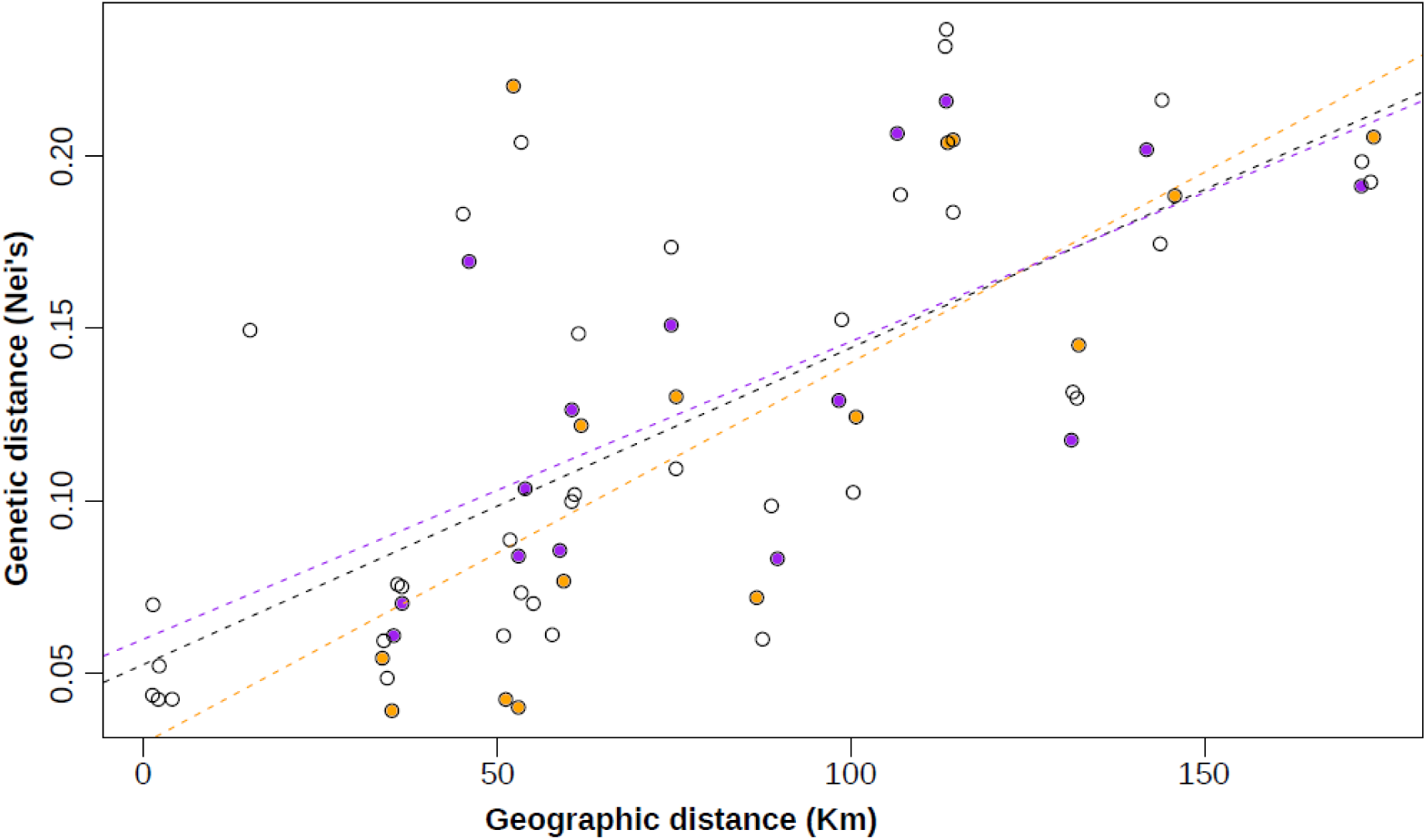
Plot of genetic vs. geographic differentiation to test for isolation-by-distance. Pairwise distances between *H. pusillum* and *H. veselskyi* populations are shown as empty circles, pairwise distances among *H. pusillum* and among *H. veselskyi* populations are shown as orange and purple filled circles, respectively. The linear regression of the pairwise distances between *H. pusillum* and *H. veselskyi* populations is shown as a black dashed line, the linear regressions for *H. pusillum* and *H. veselskyi* populations are shown as orange and purple dashed lines, respectively.

**Figure 3.**
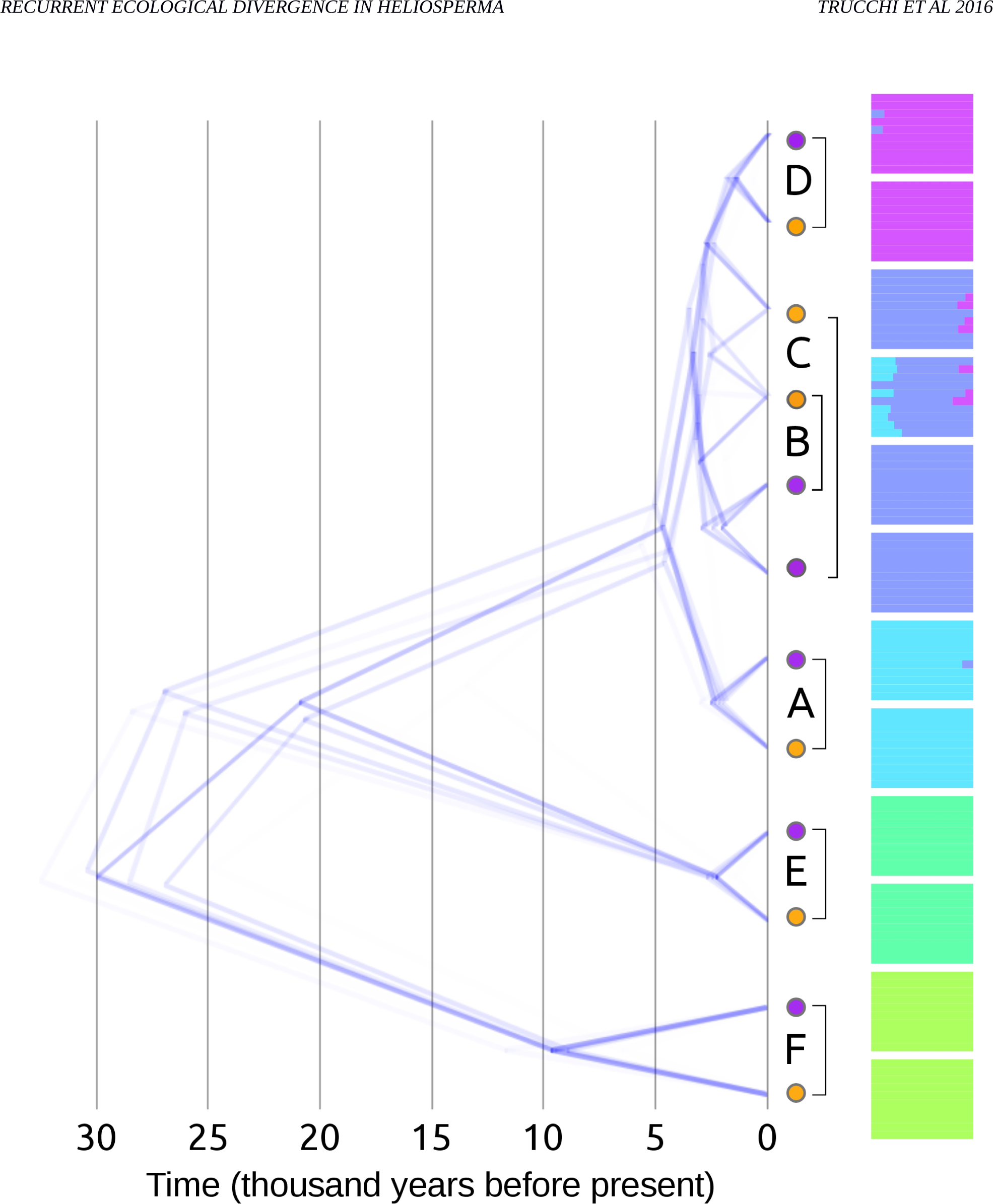
Time-calibrated phylogenetic tree and Bayesian k-means results. All alternative topologies for the population tree are shown. Barplots show inferred proportion of ancestry relative to five clusters. Colours identifying each cluster are randomly assigned and used only in this figure. *Heliosperma pusillum* and *H. veselskyi* populations are shown as orange and purple filled circles, respectively. Population labels follow Figure 1.

### Assessing the Level of Gene Flow between Species

For each population pair, the most likely number of clusters suggested by the *find.clusters* algorithm was two (Fig. 4, left panel; Fig. S7), clearly separating individuals of the two species in the *fastStructure* analysis (Fig. 4, central panel). No evidence of admixture was detected between ecotypes in D and F, whereas weak signals of admixture were uncovered in A, B, C and E. In each population pair, both reciprocal migration rates estimated with *fastsimcoal2* were below one individual per generation (Table S2). The only exception was D, where the migration rate from *H. pusillum* to *H. veselskyi* was estimated to be 1.6 individuals per generation. The two parameters and the maximum likelihood estimates were stable across replicated runs.

**Figure 4.**
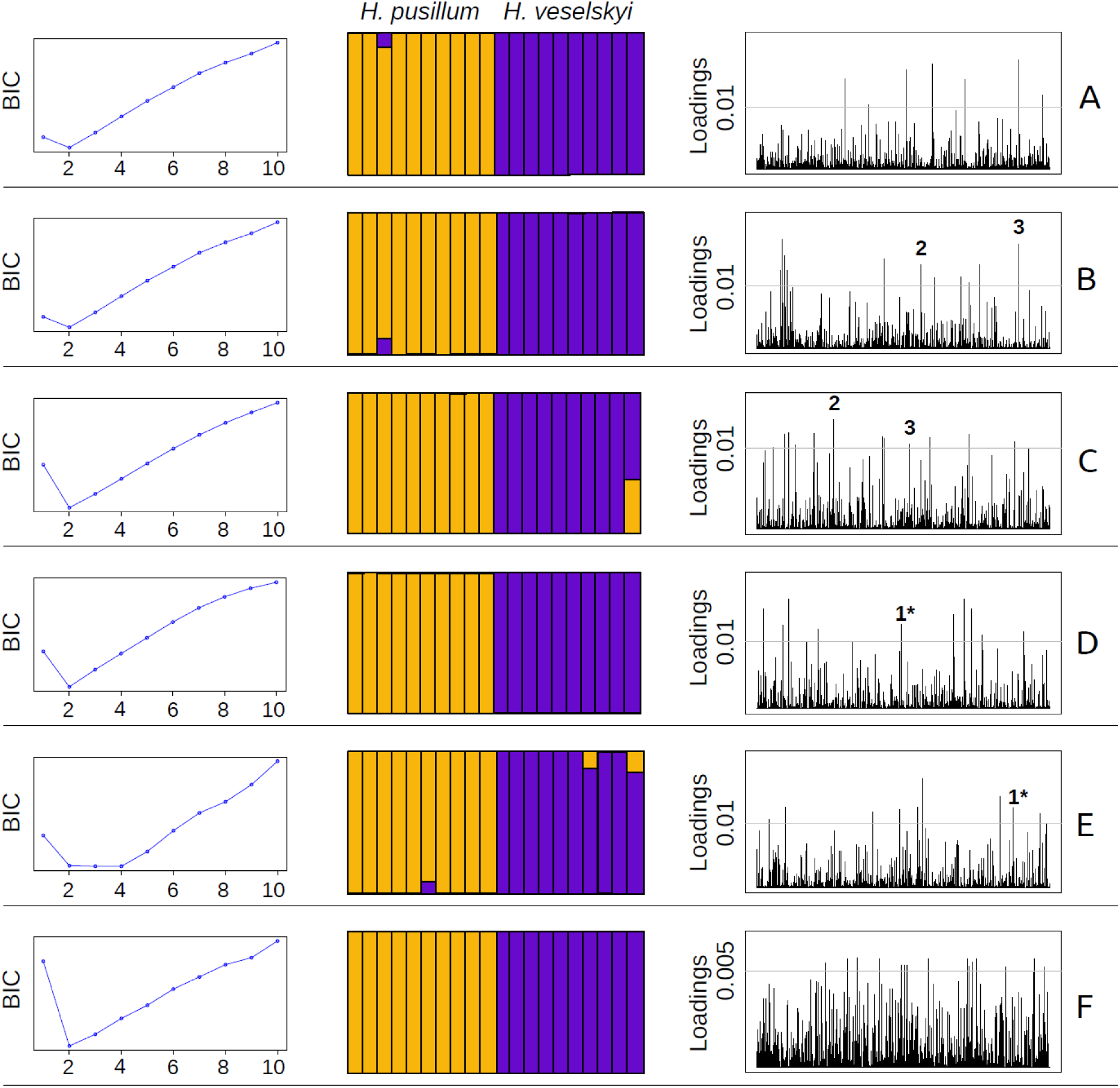
Local admixture between *Heliosperma pusillum* and *H. veselskyi*. Left panel: Bayesian Information Criterion (BIC) scores as a function of the number of clusters *(k* from 2 to 10) for successive *k*-means clustering in each population pair. Central panel: bar plots showing the ancestry of the genotypes for each population pair as inferred by the model-based Bayesian clustering for *k* = 2. Right panel: plots of the loading of each locus in Discriminant Analyses of the Principal Component in each population pair. A threshold indicating the top 1% (0.5% in F) of loci contributing to the axis explaining the separation between the species is shown as a solid horizontal line. Loci contributing in more than one population pair are indicated with numbers: 1*, same locus but alternative alleles in different population pairs; 2 and 3, same allele in different population pairs. Population labels and species colour-codes follow Figure 1.

### Detecting Outlier Loci

The SNPs that explained the divergent PCA clustering of the two species within each population pair mostly differed among population pairs (Fig. 4, right panel). The same two SNPs were contributing to the differentiation in B and C (loci 2 and 3). The same locus was found as significantly divergent in both D and E (locus 1*), but the two alleles at that SNP were differentially fixed in the two species: the allele characteristic for *H. veselskyi* in D was fixed in *H. pusillum* in E.

The haplotype-based Bayesian search for loci under selection suggested one candidate RAD locus *(locus2519;* Fig. 5), albeit with a probability only slightly exceeding the threshold of significance (logi_0_q = ‐1.3). Blasted against the published transcriptome of *Silene vulgaris*, this locus was annotated as a putative E3 ubiquitin ligase protein on *Isotig26062*. The locus featured five SNPs, but only one non-synonymous substitution showed a non-random segregation among populations and species. This SNP is in a first codon-position changing the amino acid from Glutamic acid in most *H. pusillum* individuals to Aspartic acid in most *H. veselskyi* individuals. The locus was retrieved in the RADseq data from only 43 individuals out of 120. Sanger-sequencing of this locus using primers designed out of the coding regions was used to confirm the unique amplification of this locus in 30 randomly selected individuals (Fig. S8). Primers designed within the exonic region, in contrast, produced double amplifications: sequences of *locus2519* were found together with a set of sequences of a putative paralog (Fig. S8). The paralog locus was sequenced in 30 individuals and had 12 substitutions in the 94 base pairs of the singleend RAD locus plus a 3-base pairs insertion. One substitution was in the SbfI restriction site thus preventing this paralog to be present in our RAD dataset. Nevertheless, both alleles found at this paralog locus seemed functional (no stop codon in the amplified region in the same reading frame as *locus2519)* and mapped to the same region of the *S. vulgaris* transcriptome. Orthologous alleles of *locus2519* were grouped in three major clusters: one including only *H. veselskyi* from A, B, D; one including only *H. pusillum* from B, C and F; one including *H. pusillum* from A, B, C, and D, and *H. veselskyi* from A and B. A maximum-likelihood tree of both *locus2519* and its paralog is shown in the Supplementary Material (Fig. S8).

**Figure 5.**
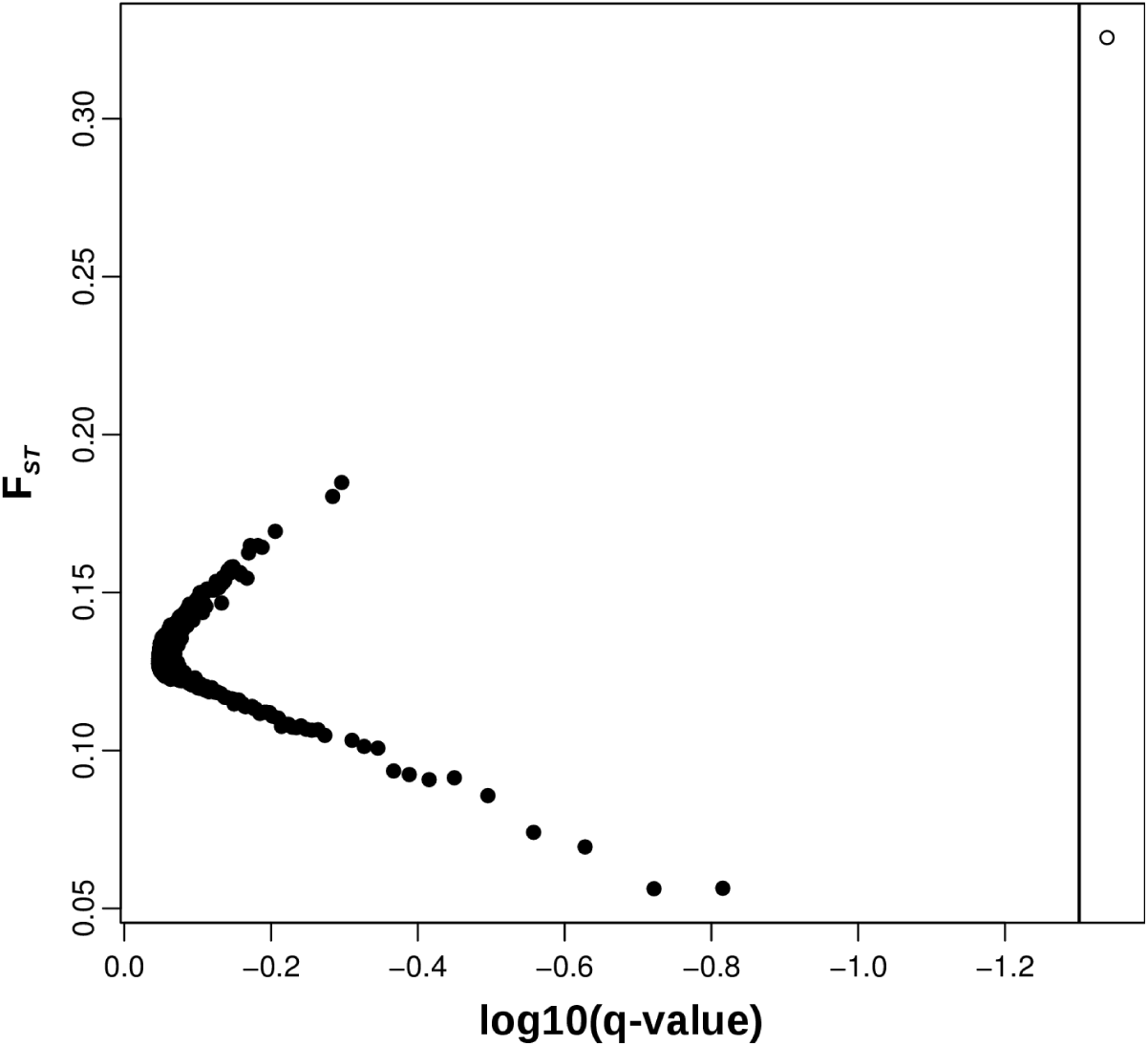
Bayesian inference of candidate loci under selection. Plot of Fst as a function of q-values for each of the 1,097 loci. A false discovery rate threshold of 0.05 is shown as vertical solid line. The candidate outlier locus *(locus2519)* is shown as an empty circle.

### Analysis of the Leaf Microbiome

The selection of loci in the RAD dataset that were restricted to either *H. pusillum* or *H. veselskyi* and found in at least one, five or eight individuals were 184,154 and 243,674, 2,301 and 2,879, and 437 and 590, respectively.

In the metagenomic analysis, only 20% of the loci were taxonomically assigned in the datasets including loci found in at least one individual, whereas between 42 and 50% were identified in the other two datasets (Fig. S9). In the dataset of loci found in at least five individuals of either species, a higher proportion of loci assigned to bacteria and fungi was found in *H. veselskyi*. Concerning the bacterial and fungal community, there were major qualitative differences between the phyllosphere of the two species (Fig. 6). In *H. pusillum* the bacterial community was characterized by the presence of Cytophagales, Sphingomonadales and Pseudomonadales, whereas Enterobacteriales and Xanthomonadales were only found in *H. veselskyi*. Bacteria from Rhizobiales and Actinomycetales were found in both species although the latter were more abundant in *H. pusillum*. Fungal taxa from Dothideomycetidae and Sordariomycetes were found only in *H. veselskyi*. Loci assigned to higher than order level (superclass Leotiomycetes) were more abundant in *H. veselskyi* (Table S3). The structure of the whole phyllosphere community was mostly consistent across all population pairs; the loci assigned to the order Xanthomonadales, in particular, co-occurred across most of the *H. veselskyi* populations (Table S3). By contrast, very few taxa were shared between the two species in each population pair (Fig. S10). A difference in the number of loci assigned to flowering plant species (summarized as Mesangiospermae) was found between the two species: 120 loci in *H. pusillum* and seven in *H. veselskyi* (Fig. S11, Table S3). Of the private loci found in *H. pusillum*, 108 mapped to plastid sequences whereas none of the seven loci private to *H. veselskyi* mapped to any plastid sequence. In addition, more than 300 loci, mainly private to four *H. pusillum* populations (Fig. S11, Table S3), were assigned to Diptera (*Drosophila*).

**Figure 6.**
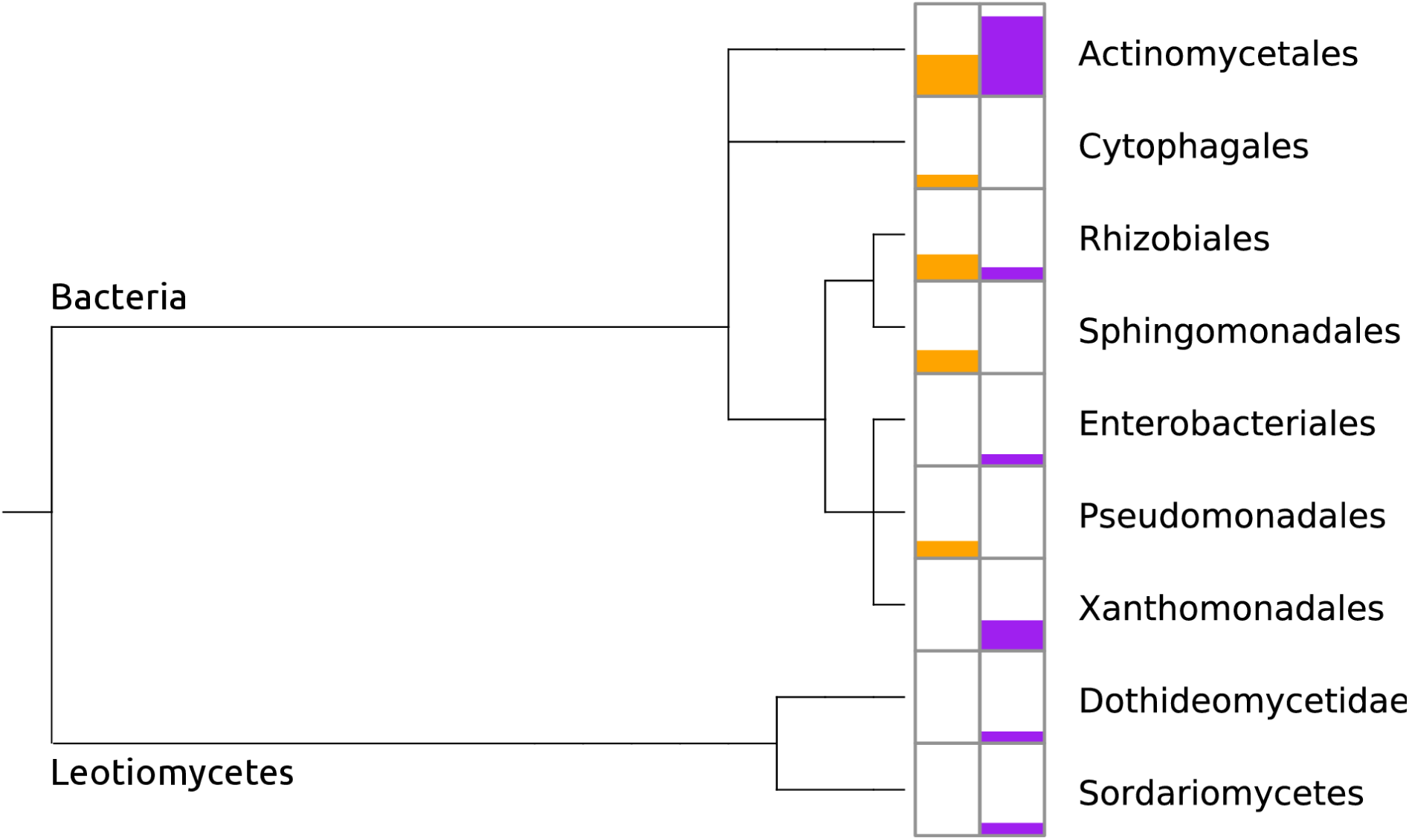
Differentiation in leaf microbiome between *H. pusillum* (shown in orange) and *H. veselskyi* (in purple). Normalized proportion of loci blasted to order level taxa of Bacteria and Leotiomycetes arranged on a phylogenetic tree downloaded from the NCBI database and analyzed in MEGAN. Detailed results of this analysis are given in the Supplementary Materials.

## Discussion

### Multiple Origins Scenario and Estimates of Gene Flow

Using genome-wide molecular data of 120 individuals of two *Heliosperma* species sampled from six population pairs over their overlapping distribution range, we surprisingly uncovered at least five independent instances of ecological divergence between the alpine *H. pusillum* and the montane *H. veselskyi*. Parallel evolution of these taxa (Figs. 3 and S4–S6) is not a singular case, as investigations of genomic data (e.g., Hohenlohe et al. 2010; Butlin et al. 2013; Roda et al. 2013; Soria-Carrasco et al. 2014) have recently reported repetitive origins in other organisms. This supports the hypothesis that recurrent origin of ecological adaptations is not restricted to hybrids and polyploids (Schluter & Nagel 1995; Levin 2001; Nosil et al. 2002; Colosimo et al. 2005), but is rather a common process in complexes that have undergone ecological diversification.

The recurrent instances of divergence took place at different localities and at different time horizons ranging roughly between 10,000 and 2,000 years before present (Fig. 3). This time range is likely an underestimate as a result of applying a genome-wide mutation rate (Ossowski *et al*. 2010) to a dataset comprising at least 41% coding loci (see section Genomic datasets from Results). The ample distribution of *H. pusillum*, from the Iberian Peninsula to the Carpathians and the southern Balkan Peninsula, as compared to the narrow range of *H. veselskyi*, restricted to the southeastern Alps, suggests the former species to be the ancestral one. A scenario of *H. veselskyi* developing from *H. pusillum* as a result of adaptation to the specific habitats is also consistent with the phylogenetic relationships within the *H. pusillum* complex, albeit based on a few loci and a limited sampling (Frajman & Oxelman 2007). We hypothesize the earliest divergence event within *Heliosperma* population pair F (Fig. 3) to have happened in unglaciated areas to the southeast of the ice sheet covering most of the Alps during the Last Glacial Maximum ca. 18,000 years ago (Fig. 1). Whereas the topology of the phylogenetic reconstruction suggests that the colonization of the Alps proceeded from East to West after the last glaciation, divergence events within the population pairs took place roughly at the same time horizon (Fig. 3) independently from the distance to the margin of the ice sheet (Fig. 1), which may serve as a proxy for the migration distance (Schönswetter et al. 2005). Divergence was likely triggered by the rapid spread of forests induced by Holocene warming (e.g., Magri et al. 2006), separating montane populations below overhanging cliffs from alpine populations. However, we cannot rule out the possibility that rare dispersal events led to the origin of, for instance, some of the constituents of population pairs B or C (Fig. 3).

Our results indicate that the populations of the two species still form a single evolutionary unit, with clear isolation by distance observed both within and across the two species (Fig. 2). This may rather relate to the limited time elapsed since the splits, than to extensive gene flow maintaining genetic contiguity. Indeed, whereas some traces of admixture or – alternatively, but difficult to disentangle – shared ancestry (i.e., incomplete lineage sorting) between the more continuously distributed *H. pusillum* populations (Poldini 2002; Wilhalm et al. 2014) are visible across the western localities A–D (Fig. 3), most of the disjunct populations of *H. veselskyi* (except B and C) seem today isolated from each other. This is consistent with the small population size observed in the *H. veselskyi* localities, rendering faster genetic drift and accumulation of some classes of TEs (Supplementary Information §2 and Figs. S1–S2) likely.

Within each population pair, interspecific gene flow (Table S2) and admixture (Fig. 4) are minimal, even over distances of less than two kilometres. Growing sites of *H. veselskyi*, which are usually situated below overhanging rocks in the forest belt, may be difficult to reach for the dispersed seeds of *H. pusillum*. A further, not mutually exclusive, hypothesis is that traits of one species are maladaptive at the growing sites of the other species as predicted by models of ecological speciation (Schluter 2009). Transplantation experiments are currently being performed to better understand this aspect. Further, it is unclear if ecological segregation of pollinators contributes to reproductive isolation (Lowry et al. 2008; Whitehead & Peakall 2014). Selection against hybrids (Orr 1995) as well as reproductive isolation caused by BDMI have been identified in animals and plants (Presgraves 2010), causing double heterozygote individuals to be inviable or have lower fitness under certain ecological conditions. The BDMI model can be extended to many loci, whose complex epistatic interaction may lead to hybrid fitness reduction or inviability (Bomblies & Weigel 2007) and a very rapid evolution of reproductive barriers (Skrede et al. 2008). Interestingly, hybrid weakness may result from BDMI occurring at immunity-related genes (Bomblies & Weigel 2007; Rieseberg & Willis 2007), potentially associated with different pathogens load. Crossing experiments between *H. pusillum* and *H. veselskyi* in a common garden did not show any clear sign of reduced viability or fertility of hybrids across two generations (Bertel et al., in press), indicating once again that this system is at an early stage in the speciation process. However, intrinsic hybrid weakness may appear at a later generation (e.g., Chapman et al. 2015) and/or hybrids might be selected against only in the natural conditions of the parental habitats.

*Heliosperma veselskyi* populations are likely to proceed in the speciation process and become fully isolated both from *H. pusillum* and from each other (see Lowry 2012). A phylogenetic analysis of the *H. pusillum* group (Frajman & Oxelman 2007) suggests that the process unravelled in the Alps could have been common in the diversification history of this species complex and that the low-elevation endemics on the Balkan Peninsula could have similarly evolved via ecological speciation. Polyphyletic phylogenies (Hörandl 2006) and the presence of cryptic species (e.g., Skrede et al. 2008) are likely outcomes of such complex evolutionary process observed in the Alps at an early stage.

### Adaptive Loci Search

The similar morphological and ecological adaptation across different *H. veselskyi* populations could be the result of similar selective pressures acting on a largely common pool of genetic diversity (i.e. collateral evolution by shared ancestry; Schluter 2009; Stankosky 2013; Stern 2013). Only one out of 1,097 variable loci screened here reported a signature of non-random segregation among populations of the two species (Figs. 5, S8). This locus has been annotated as an E3 ubiquitin ligase protein, a large family of enzymes that also plays a role in plant innate immune system (see Duplan & Rivas 2014 for a review), which is of interest given the biotic differentiation at the level of the phyllosphere between the two species (Fig. 6). A lack of similar genomic divergence at the same traits across the six replicated population pairs, which show instead similar morphological and ecological divergence, has been reported in other biological systems as well (e.g., Ravinet et al. 2015 and references therein). Independent and different mutations occurring in the same gene, or polygenic traits underlying the ecological adaptations, could explain these results. However, given the very short divergence time of each *Heliosperma* population pair (roughly less than 10,000 years, Fig. 3), a significant contribution of *ex novo* mutations can be ruled out and repetitive origins from standing variation appear more likely (Barrett & Schluter 2008). On the other hand, as adaptive traits are likely polygenic (Pritchard et al. 2010), the signature of selection may be difficult to detect unless more advanced analytical approaches, not applicable in our case, are employed (Daub et al. 2013). The reduced representation RADseq dataset employed here screened only a limited portion of the genome (0.1%) and of the genes (ca 3.5% – assuming ca. 20,000 genes in the *Heliosperma* genome and ca. 700 RAD loci mapping to the *Silene vulgaris* transcriptome and gene sequences in GenBank). It is thus possible that the adaptive loci, or linked neutrally-evolving loci, were not covered in our genome scan. In addition, if selection is acting on standing genetic variation present in the ancestral population or if very large effective population sizes characterize one or both diverging populations (Pritchard et al. 2010), the signature of divergence around loci under selection is expected to be minimal (Hermisson & Pennings 2005). These aspects could then exacerbate the difficulties in finding adaptive loci that, in some cases, have been shown to be restricted to one single SNP (e.g., O’Brown et al. 2015).

Morphological and ecological divergence shared by replicated instances of adaptation can also be due to similar changes in gene expression under epigenetic control (e.g., Bossdorf & Zhang 2011; Latzel *et al*. 2013). Indeed, epigenetic changes can alter the expression of genes (e.g., Rapp & Wendel 2005; Li *et al*. 2012; Le *et al*. 2015), and can appear at a very fast rate and recurrently at certain genomic positions (Becker *et al*. 2011; Smitz *et al*. 2011). These aspects could then be particularly important at the early stages of population divergence and, in particular, in the case of repeated ecological adaptation. A dedicated study on the divergence of cytosine methylation profiles employing the same samples as in the experiment presented here is currently ongoing.

### Characterizing the Leaf Microbiome using RADseq Data

We report here evidence of a clear similarity in the structure and diversity of the phyllosphere community across populations of each species (Table S3) contrasted by a marked differentiation between the species (Figs. 6, S9–S11). Each species-specific assemblage of the phyllosphere community is likely driven by the ecological, morphological and physiological divergence (Bertel *et al*., in press) already in place between the two species inhabiting wet alpine and dry montane habitats, respectively. Nonetheless, the different sets of biotic interactions can, indeed, reinforce this divergence. Bacterial and fungal organisms, including commensal, mutualistic and pathogenic taxa living on roots and leaves, have been shown to have a deep physiological, functional and evolutionary impact on the host plant species (e.g., Guttman *et al*. 2014; Kembel *et al*. 2014; Lebeis 2015) and, consequently, on ecological adaptation and evolution (Margulis & Fester 1991; Zilber-Rosenberg & Rosenberg 2008). Including the analysis of the microbiome in molecular ecology studies has been identified as one of the main priorities in the field (Andrew *et al*. 2013), especially through the serendipitous analysis of microbiome-associated contamination of host genome-wide sequencing (Kumar & Blaxter 2011). Here, we demonstrate that a high-coverage RADseq dataset can serve as preliminary screen of the phyllosphere of target species. Follow-up investigations should focus on *i*) the identification of microbial taxa in the phyllosphere of *H. pusillum* and *H. veselskyi* through in-depth metagenomic analyses, *ii*) the characterization of *Heliosperma*-specific microbiomes (both in phyllosphere and rhizosphere) contrasted with the microbiomes of accompanying plant species, thereby allowing for disentangling the habitat-specific microbial background from *Heliosperma*-specific taxa, and *iii*) the characterization of the functional aspects of the most important biotic interactions.

## Acknowledgements

We would like to remember here Ruth Flatscher that initially was involved in this study, but sadly passed away. Maria Lorenzo Romero is acknowledged for constructing some of the RADseq libraries. This work was supported by the Austrian Climate Research Programme (ACRP5-EpiChange-KR12AC5K01286 to P.S.); and the Austrian Science Fund (FWF, Y661-B16 to O.P.).

## Supplementary Information

Data and additional results available from the Dryad Digital Repository (doi:XXXX). Raw genomic data available from the NCBI Short Reads Archive (acc. num. SRP065672, SRP068291).

## Genomic and Metagenomic Analyses Reveal Parallel Ecological Divergence in *Heliosperma pusillum* (Caryophyllaceae)

Emiliano Trucchi, Božo Frajman, Thomas HA Haverkamp, Peter Schönswetter, Ovidiu Paun

## Supplementary Informations

## 1. Sampling information

**Table S1. Sample design** (labels as in Fig. 1 in the main text).

**Population, Species, Bioproject Accession, Sample ID, SRA accession, Locality, Lat_Lon, Altitude, Collected by, Collection date**

**A**, *H. pusillum*, PRJNA308183,PRJNA300879, PVA3-PVA4-PVA5-PVA6-PVA7-PVA8-PVA9-PVA11-PVA14-PVA18, SRR2891253-SRR2891254-SRR2891255-SRR2891235-SRR3096515-SRR2891236-SRR2891249-SRR2891250-SRR2891251-SRR2891252, Italy: Trentino-Alto Adige: Dolomiti di Gardena/Grödner Dolomiten, 46.601 N 11.768 E, 2290, Ruth Flatscher, 24-Jul-2011

**A**, *H. veselskyi*, PRJNA308183,PRJNA300879, VVA4-VVA6-VVA12-VVA15-VVA16-VVA19-VVA20-VVA23-VVA26-VVA29, SRR2891237-SRR2891238-SRR2891239-SRR2891243-SRR2891244-SRR2891245-SRR2891246-SRR2891247-SRR3096516-SRR2891256, Italy: Trentino-Alto Adige: Dolomiti di Gardena/Grödner Dolomiten, 46.564 N 11.77 E, 1690, Ruth Flatscher, 24-Jul-2011

**B**, *H. pusillum*, PRJNA308183, PTO5-PTO6-PTO8-PTO12-PTO19-PTO2-PTO20-PTO22-PTO25-PTO27, SRR3096551-SRR3096552-SRR3096557-SRR3096553-SRR3096554-SRR3096555-SRR3096556-SRR3096558-SRR3096559-SRR3096560, Italy: Trentino-Alto Adige: Dolomiti di Braies/ Pragser Dolomiten,, 46.644 N 12.205 E, 2190, Ruth Flatscher, 8-Jul-2011

**B**, *H. veselskyi*, PRJNA308183, VTO8-VTO10-VTO11-VTO14-VTO16-VTO21-VTO23-VTO27-VTO28-VTO29, SRR3096614-SRR3096615-SRR3096616-SRR3096617-SRR3096618-SRR3096619-SRR3096620-SRR3096621-SRR3096622-SRR3096623, Italy: Trentino-Alto Adige: Dolomiti di Braies/ Pragser Dolomiten,, 46.645 N 12.233 E, 1420, Ruth Flatscher, 8-Jul-2011

**C**, *H. pusillum*, PRJNA308183, Pa15-PCI16-Pa17-Pa18-Pa19-Pa20-Pa22-Pa23-Pa24-Pa30, SRR3096512-SRR3096513-SRR3096527-SRR3096538-SRR3096550-SRR3096561-SRR3096572-SRR3096583-SRR3096601-SRR3096613, Italy: Friuli-Venezia Giulia: Val Cimoliana, 46.391 N 12.48 E, 1700, Ruth Flatscher, 7-Jul-2011

**C**, *H. veselskyi*, PRJNA308183, Va12-VCI15-Va20-Va21-Va22-Va23-Va24-Va26-Va27-VaC, SRR3096562-SRR3096563-SRR3096564-SRR3096565-SRR3096566-SRR3096567-SRR3096568-SRR3096569-SRR3096570-SRR3096571, Italy: Friuli-Venezia Giulia: Val Cimoliana, 46.38 N 12.489 E, 1180, Ruth Flatscher, 7-Jul-2011

**D**, *H. pusillum*, PRJNA308183, PHO5-PHO7-PHO11-PHO13-PHO15-PHO16-PHO2-PHO22-PHO23-PHO30, SRR3096528-SRR3096529-SRR3096530-SRR3096531-SRR3096534-SRR3096532-SRR3096533-SRR3096535-SRR3096536-SRR3096537, Austria: Kärnten: Lienzer Dolomiten, 46.762 N 12.877 E, 2055, Ruth Flatscher, 3-Aug-2011

**D**, *H. veselskyi*, PRJNA308183, VHO6-VHO11-VHO13-VHO16-VHO17-VHO18-VHO19-VHO21-VHO25-VHO29, SRR3096584-SRR3096585-SRR3096587-SRR3096593-SRR3096595-SRR3096596-SRR3096597-SRR3096598-SRR3096599-SRR3096600, Austria: Kärnten: Lienzer Dolomiten, 46.774 N 12.901 E, 790, Ruth Flatscher, 3-Aug-2011

**E**, *H. pusillum*, PRJNA308183, PNE6-PNE11-PNE14-PNE15-PNE16-PNE19-PNE21-PNE25-PNE26-PNE28, SRR3096539-SRR3096540-SRR3096541-SRR3096542-SRR3096543-SRR3096544-SRR3096546-SRR3096547-SRR3096548-SRR3096549, Italy: Friuli-Venezia Giulia: Alpi Giulie, 46.376 N 13.459 E, 1820, Ruth Flatscher, 6-Jul-2011

**E**, *H. veselskyi*, PRJNA308183, VNE2-VNE20-VNE22-VNE24-VNE25-VNE5-VNE67-VNE73-VNE74-VNE81, SRR3096606-SRR3096602-SRR3096603-SRR3096604-SRR3096605-SRR3096607-SRR3096608-SRR3096609-SRR3096610-SRR3096611, Italy: Friuli-Venezia Giulia: Alpi Giulie, 46.388 N 13.459 E, 1170, Ruth Flatscher, 6-Jul-2011

**F**, *H. pusillum*, PRJNA308183, PDI10-PDI16-PDI17-PDI19-PDI20-PDI21-PDI24-PDI25-PDI26-PDI7, SRR3096514-SRR3096517-SRR3096518-SRR3096519-SRR3096520-SRR3096521-SRR3096522-SRR3096523-SRR3096524-SRR3096526, Slovenia: Primorska: Trnovski gozd, 45.989 N 13.845 E, 1100, Božo Frajman, 17-Jul-2011

**F**, *H. veselskyi*, PRJNA308183, VDI4-VDI8-VDI11-VDI25-VDI27-VDI29-VDI3-VDI30-VDIM3-VDIM7, SRR3096573-SRR3096574-SRR3096575-SRR3096576-SRR3096578-SRR3096577-SRR3096579-SRR3096580-SRR3096581-SRR3096582, Slovenia: Primorska: Idrijca valley, 46.117 N 13.911 E, 540, Božo Frajman, 17-Jul-2011

## 2. Estimating the proportion of transposable elements in the Heliosperma RADseq dataset

In order to detect significant differences in TE abundance between populations of *H. veselskyi* and *H. pusillum* we used all RADseq paired-end reads of each individual and the database of transposable elements identified in plants and collected in the RepBase database on the Giri repository (http://www.girinst.org/). We downloaded as a plain fasta file all elements matching “Viridiplantae” in the database, including transposable elements, simple repeats, pseudogenes and integrated viruses. We then indexed this file and separately blasted to this reference all paired-end reads for each individual sample using a maximum e-value of 0.0001. For each individual, we estimated the proportion of reads with successful hit to the TE database. In each population pair, a Kolmogorov-Smirnov test *(R* function *ks.test* in the *stats* package) was implemented to test for significant differentiation in the proportion of TEs. We then grouped the successful hits by TE family and counted the occurrence of each family in each individual. We then selected 11 families of TEs that were found in each of the 120 individuals. After grouping the individuals by species, we assessed the difference in the presence of each family between the two species, again with a Kolmogorov-Smirnov test.

The percentage of paired-end reads mapping to the Viridiplantae TE database (Fig. S1) was in general low for all individuals spanning from 0.46% (21,757 hits) to 4.1% (45,489 hits). At the level of population pairs a significantly higher proportion of TE hits in *H. veselskyi* as compared to *H. pusillum* was found in HO and DI (*p* < 0.05), and, marginally significant, in TO (*p* = 0.06). In VA and NE, the distribution of hits to individuals of the two species was very similar, whereas *H. pusillum* in DI showed a higher proportion of TEs than *H. veselskyi* (Fig. S1). Across all six population pairs more individuals with higher proportion of hits to transposable elements were found in *H. veselskyi*. However, when grouping all individuals by species (Fig. S2), the difference in the distributions of reads mapping to TEs was barely significant (*p*-value = 0.047). Gypsy and Copia (Class I Retrotransposons, order LTR) were the most represented superfamilies and more abundant in *H. veselskyi*, although the difference was not significant (Fig. S2). Retrotransposons belonging to the order LINE, superfamily L1, appeared to be significantly more represented in *H. veselskyi*. In the Class II DNA Transposons, the superfamily CATCA, order TIR, was significantly more abundant in *H. veselskyi*. On the contrary, the family SINE2 of the SINE order was more abundant in *H. pusillum*.

Our results exclude any significant bias in our dataset due to RAD loci representing TEs. As our RADseq dataset was enriched for genes (42% of the loci) and genomic clusters of genes are generally poor in TEs (Schmidt and Heslop-Harrison 1998), the low proportion of reads mapping to TEs is not surprising. Nevertheless, we found a slightly higher proportion of transposable elements in three out of six populations of *H. veselskyi* and a general increase in the amount of retrotransposons, especially Gypsy and Copia, in this species. As expected, these two retrotransposon families were the most represented in our dataset thus driving the global pattern. Also L1, LINE, elements appear to be more represented in *H. veselskyi*. Among Class II DNA Transposons, CATCA elements are more abundant in *H. veselskyi*. This family of TEs has been shown to be involved in intron/exon re-arrangements (i.e., alternative splicing) and in changing regulatory pathways of genes, thus likely being important in the early phases of adaptation (Zabala and Vodkin 2007, Alix et al 2008, Buchmann et al 2014). Being just a by-catch of our RADseq experiment, our screening is only superficially describing TE patterns but it highlights the interest in further investigations focused on TEs and on the role they may play in different stages of populations divergence.

---

**Figure S1.**
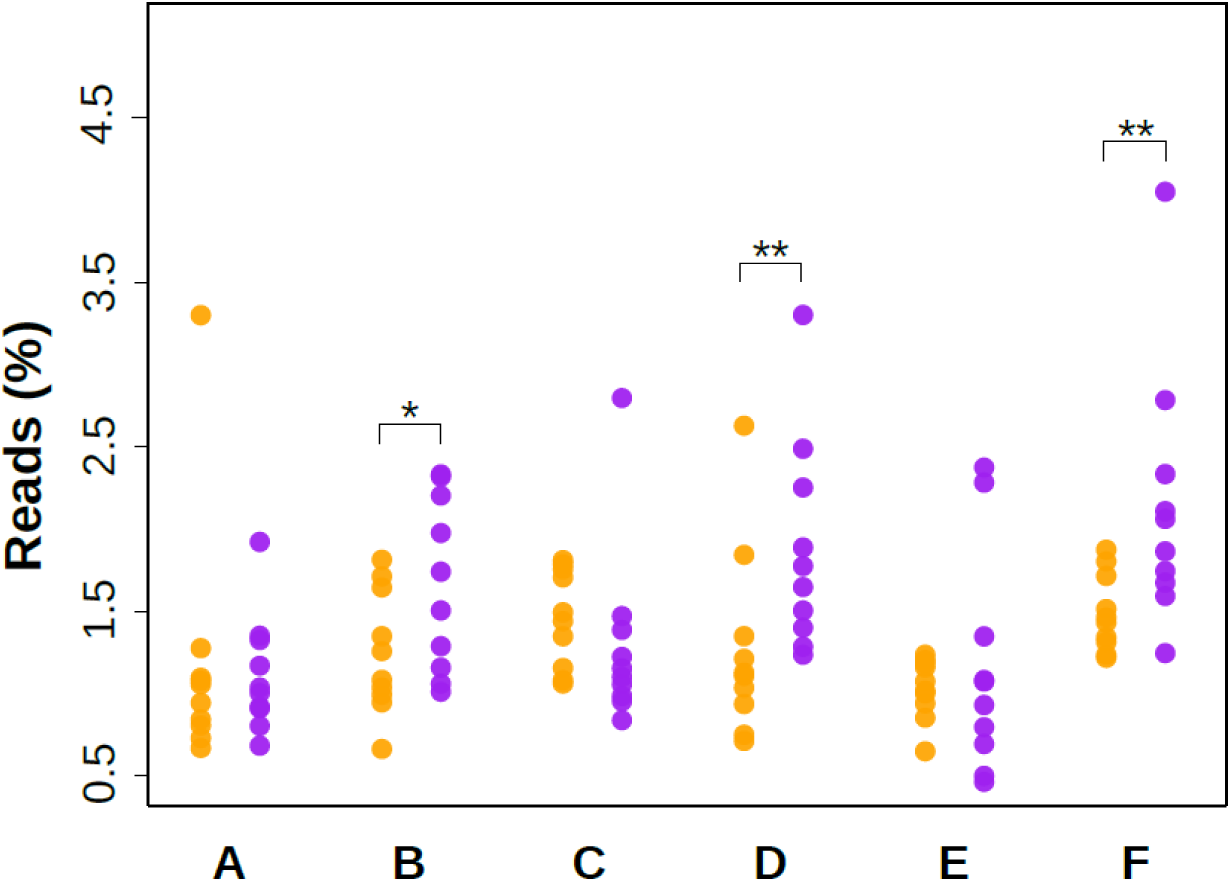
Proportion of transposable elements in 120 individuals of *Heliosperma pusillum* (orange) and *H. veselskyi* (purple) sampled from six population pairs. Illustrated are percentages of paired-end reads showing significant (blast e-value < 0.0001) hits to any element listed in the *Giri* database of plant transposable elements. Significant differentiation in the distribution of individual hits between species is indicated: * *p*-value = 0.06; ** *p*-value < 0.05. See main text for population labels.

**Figure S2.**
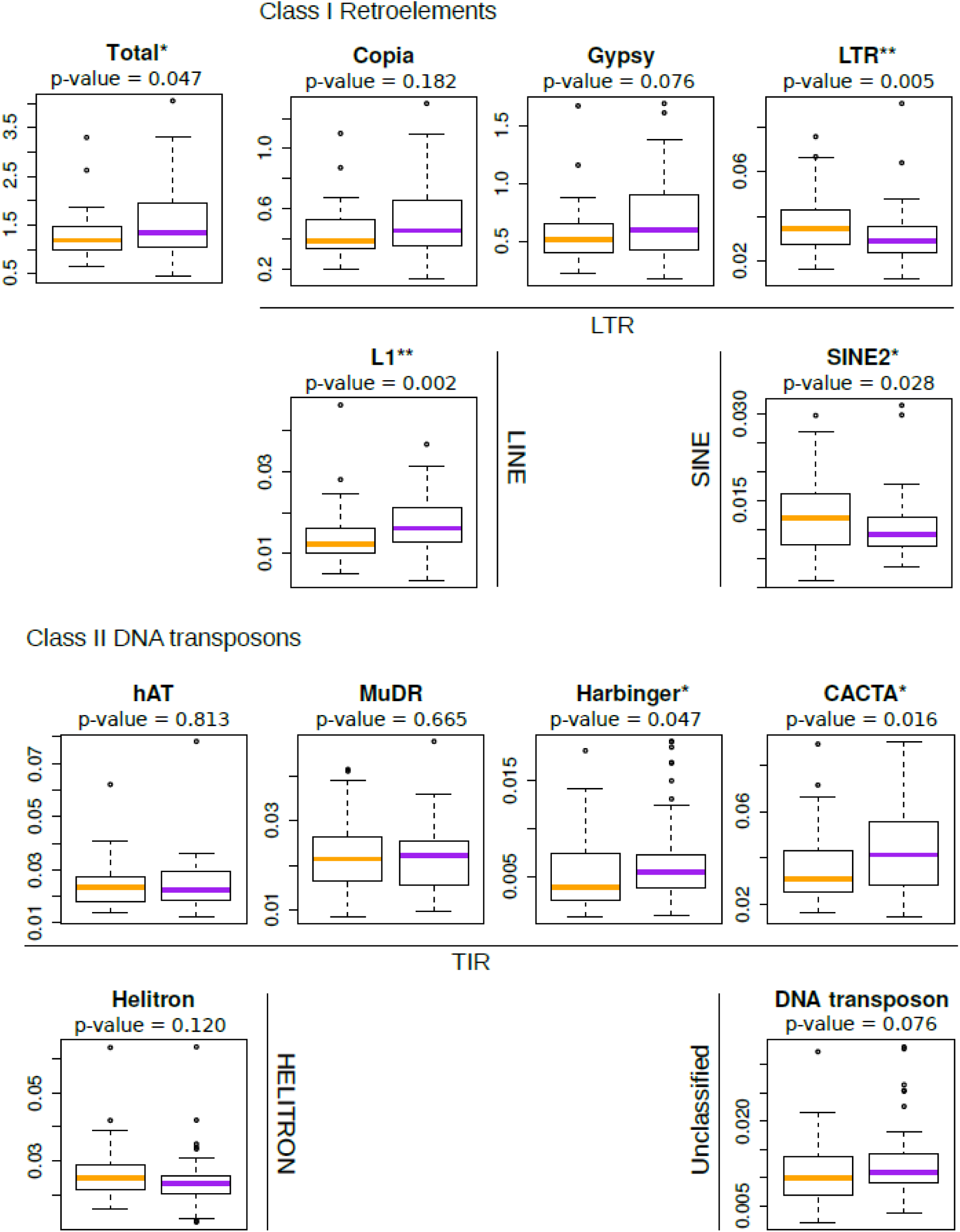
Species-specific proportion of transposable elements in *Heliosperma pusillum* (orange) and *H. veselskyi* (purple). Boxplots show the percentage of paired-end reads of each individual mapping to plant transposable elements. Superfamilies of transposable elements are grouped in Orders (LTR, LINE, and SINE, and TIR, HELITRON, and Unclassified) within the respective Class (I Retrotransposons and II DNA Transposons). P-values according to Kolmogorov-Smirnov tests are indicated: \**p*-value < 0.05; ** *p*-value < 0.01.

## 3. Dataset attributes and removal of bacterial contamination

**Figure S3.**
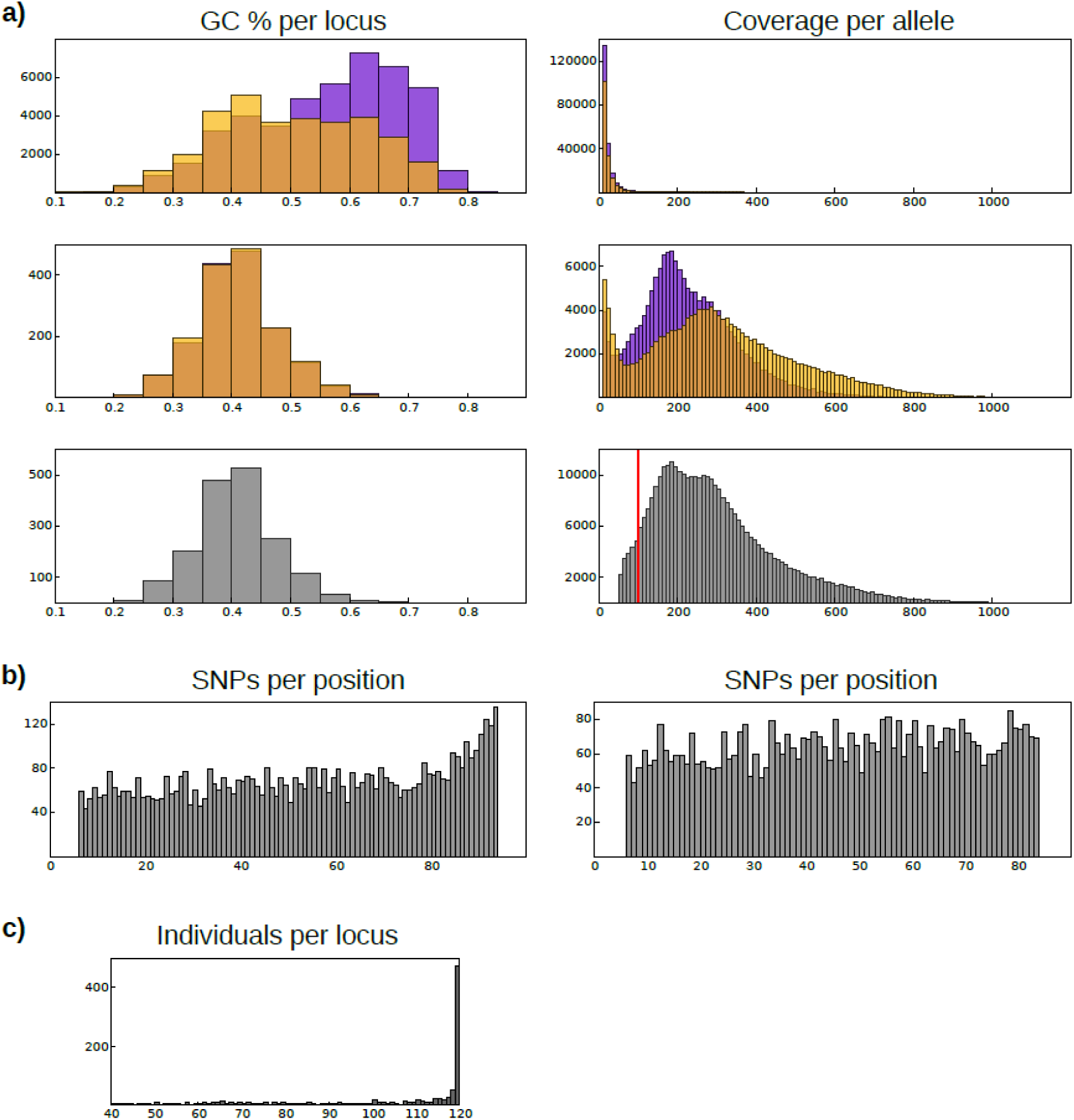
Dataset quality control. a) GC content per locus (left) and coverage per allele (right). Top panel: loci sequenced in two to five individuals; middle panel: loci sequenced in 30 to 60 individuals; bottom panel: loci sequenced in 40 to 120 individuals. Loci assembled in catalogues restricted to *H. pusillum* and *H. veselskyi* are shown in orange and purple, respectively; loci assembled in the final catalog including both species are in grey. Count of loci is on the y-axis. The threshold for the minimum coverage necessary to call a stacks (-m 100) in the final dataset is shown by a vertical red line. Note that homozygous loci are counted as two alleles and their coverage is divided by two so that alleles showing a coverage lower than the threshold are present in the plot. The difference in the distribution is not due to a library effect as the two species were sequenced in mixed libraries. b) Distribution of SNPs along the locus length before filtering (left panel) and in the final dataset (right panel) including both species. Count of SNPs is on the y-axis. c) Number of individuals sequenced per locus in the final dataset. Count of loci is on the y-axis.

## 4. Data filtering

The custom python script (*loci_selector_v2.py*) employed for filtering the output file generated by the function *export_sql.pl* in the *Stacks* package (see main text for details) is freely available for download from **http://www.emilianotrucchi.it/images/loci_selector_v2.py.**

## 5. Testing the hypothesis of parallel evolution

**Figure S4.**
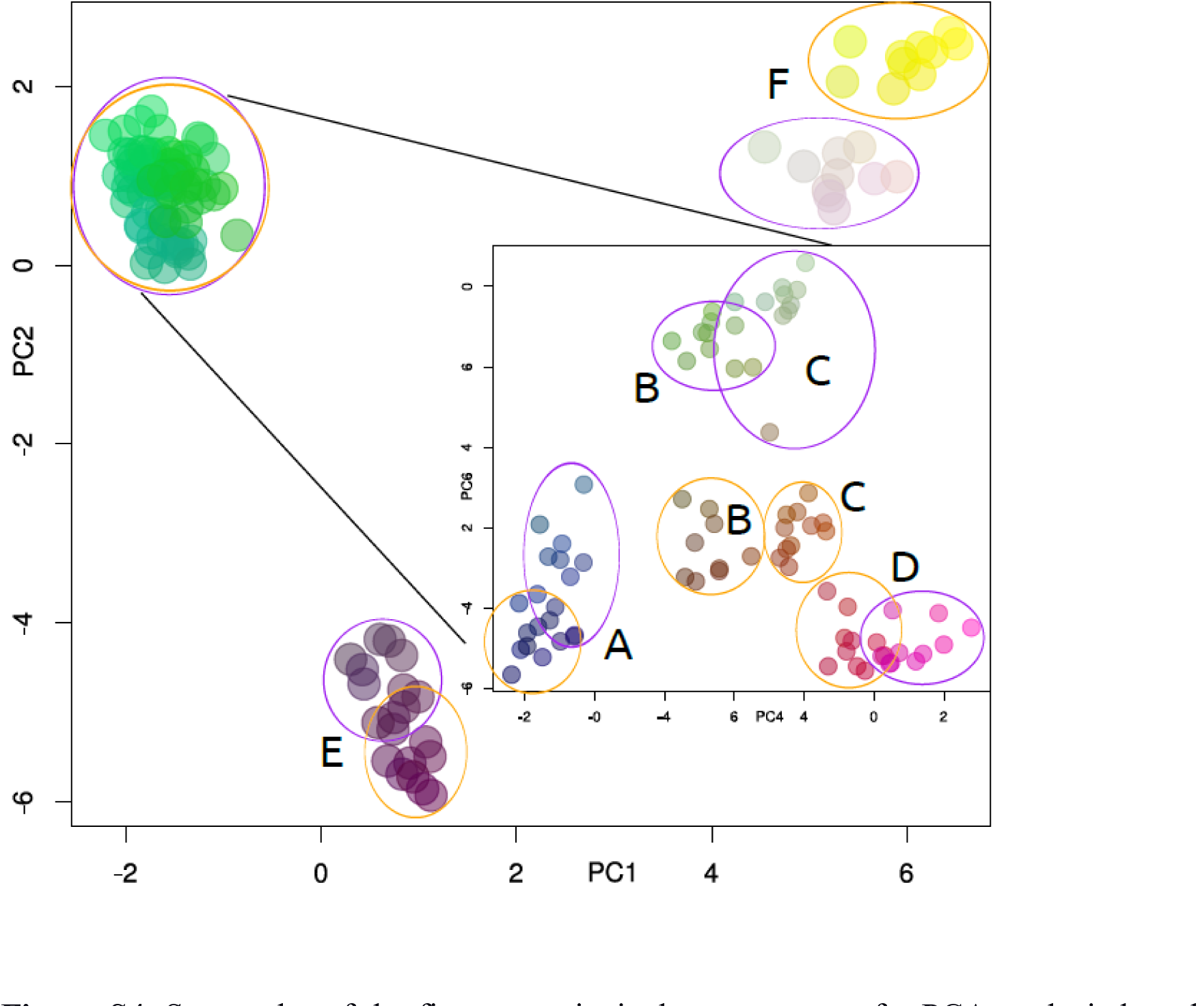
Scatterplot of the first two principal components of a PCA analysis based on 1,097 loci from 120 individuals of *Heliosperma pusillum* and *H. veselskyi*. The circle filling colours were obtained using the first three principal components as separate RGB channels. In the inset, a PCA performed on 80 individuals from population pairs A–D is shown. The colour of the ellipses reflects taxonomy (orange, *Heliosperma pusillum*; purple, *H. veselskyi)*. Population labels are as in Figure 1 in the main text.

**Figure S5.**
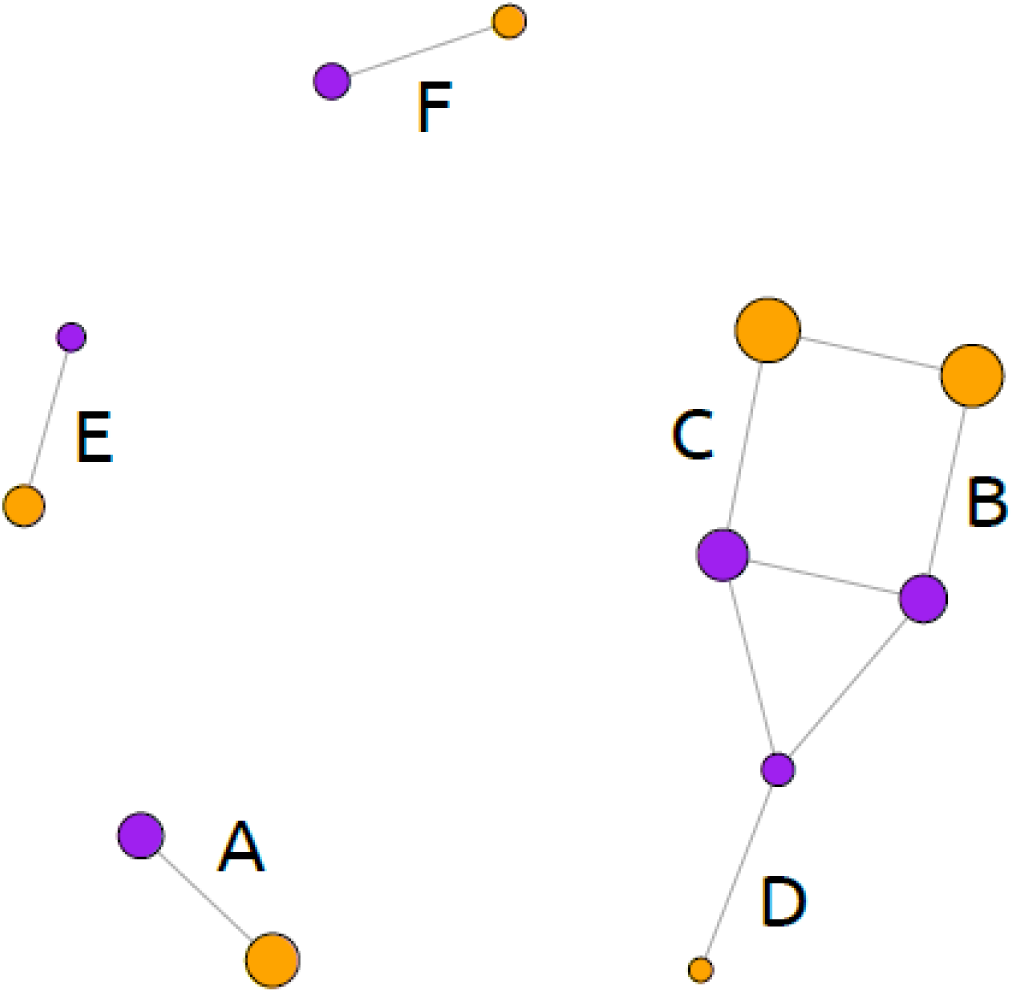
Covariance structure among 12 populations of *Heliosperma pusillum* (orange) and *H. veselskyi* (purple) forming six population pairs. Shown is a population network based on the conditional graph distance as inferred in *popGraph* (see main text for details and reference). Edges are drawn among populations showing co-variance exceeding average co-average. The size of each circle is proportional to population variance. Population labels are as in Figure 1 in the main text.

**Figure S6.**
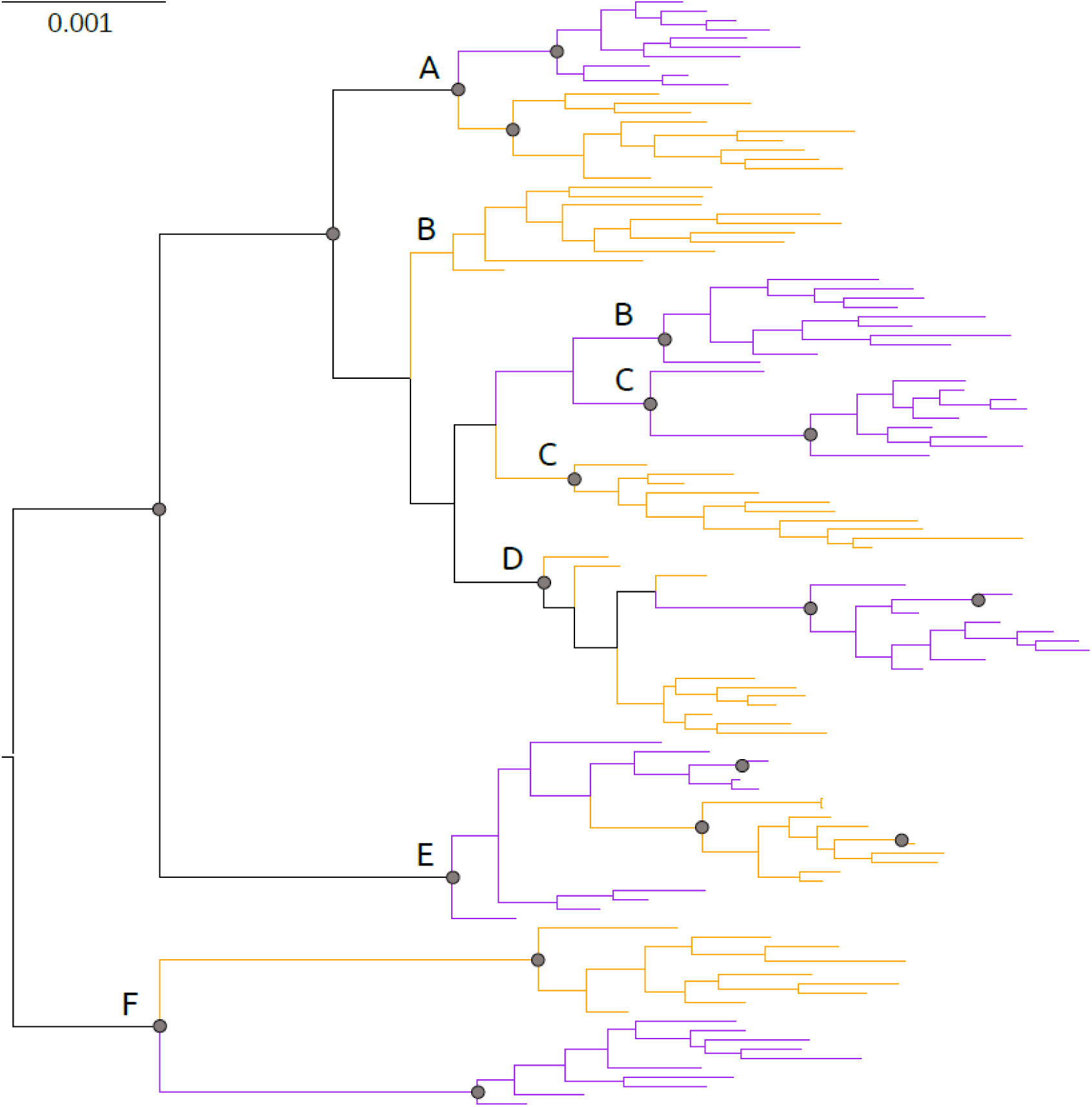
Maximum-Likelihood phylogenetic tree inferred by RaxML (see main text for details) illustrating relationships among 120 individuals of *Heliosperma pusillum* (orange) and *H. veselskyi* (purple). The mid-point rooted tree is based on 1,097 concatenated loci coding heterozygous sites as IUPAC ambiguities. Bootstrap values > 95 are represented as grey circles. Population labels are as in Figure 1 in the main text.

## 6. Assessing the level of gene flow between species

**Table S2.** Mean values of asymmetric migration rates in each population pair of *H. pusillum* and *H. veselskyi* and maximum composite-likelihood (Max L) of the underlying isolation-with-migration model as inferred in *fastsimcoal2* (see main text for details): m P->V, migration from *H. pusillum* to *H. veselskyi*; m V->P, migration from *H. veselskyi* to *H. pusillum*. Population labels are as in Figure 1.

|  | m P->V | m V->P | Max L |
| --- | --- | --- | --- |
| A | 0.95 | 0.53 | -2282 |
| B | 1.63 | 0.10 | -2497 |
| C | 0.52 | 0.06 | -1964 |
| D | 0.66 | 0.06 | -1733 |
| E | 0.44 | 0.07 | -2054 |
| F | 0.12 | 0.08 | -2961 |

**Figure S7.**
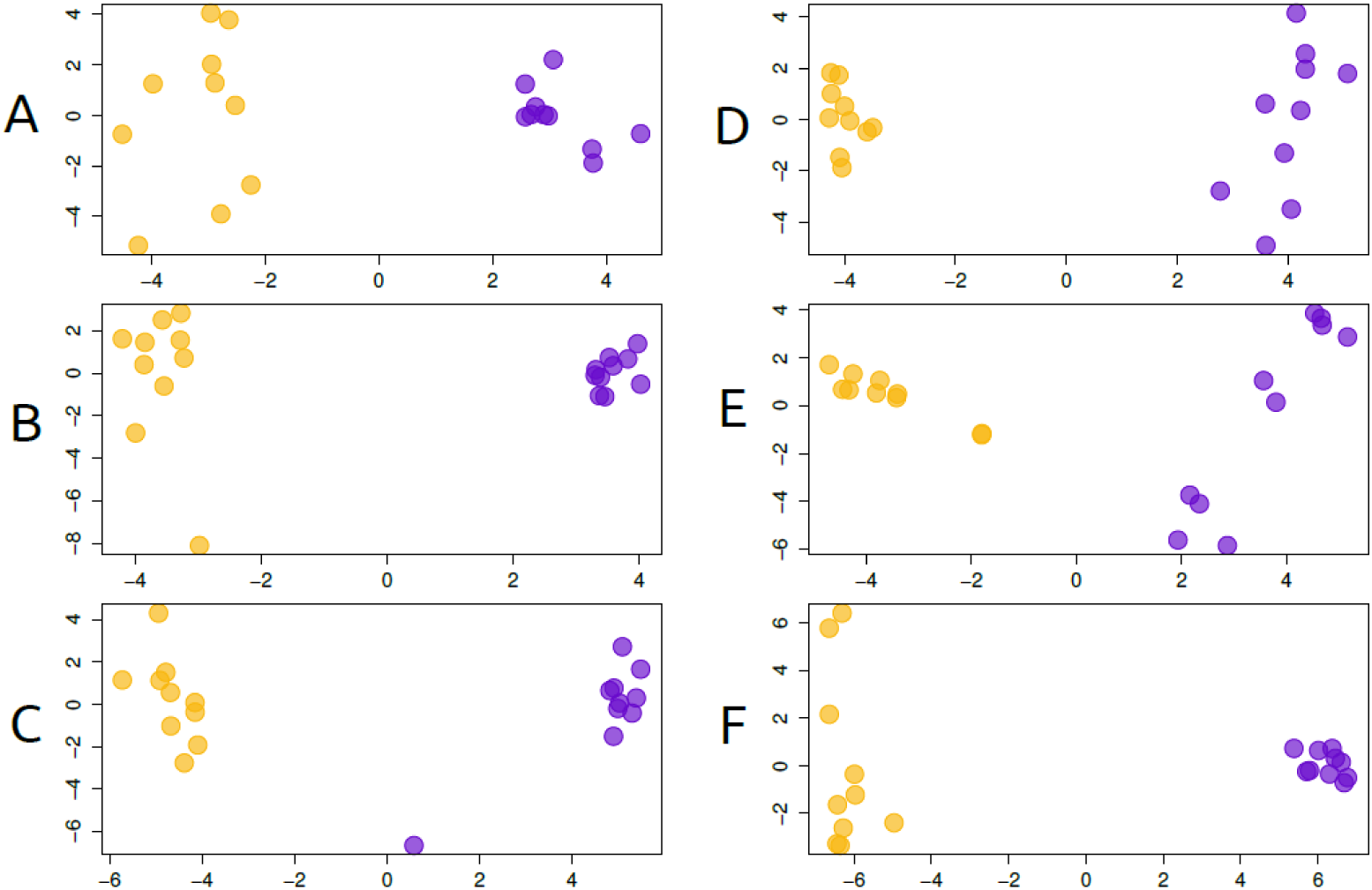
Admixture between *Heliosperma pusillum* (orange) and *H. veselskyi* (purple). Shown are scatterplots of the first two components of a PCA of each population pair. Population labels are as in Figure 1 in the main text.

## 7. Detecting outlier loci

**Figure S8.**
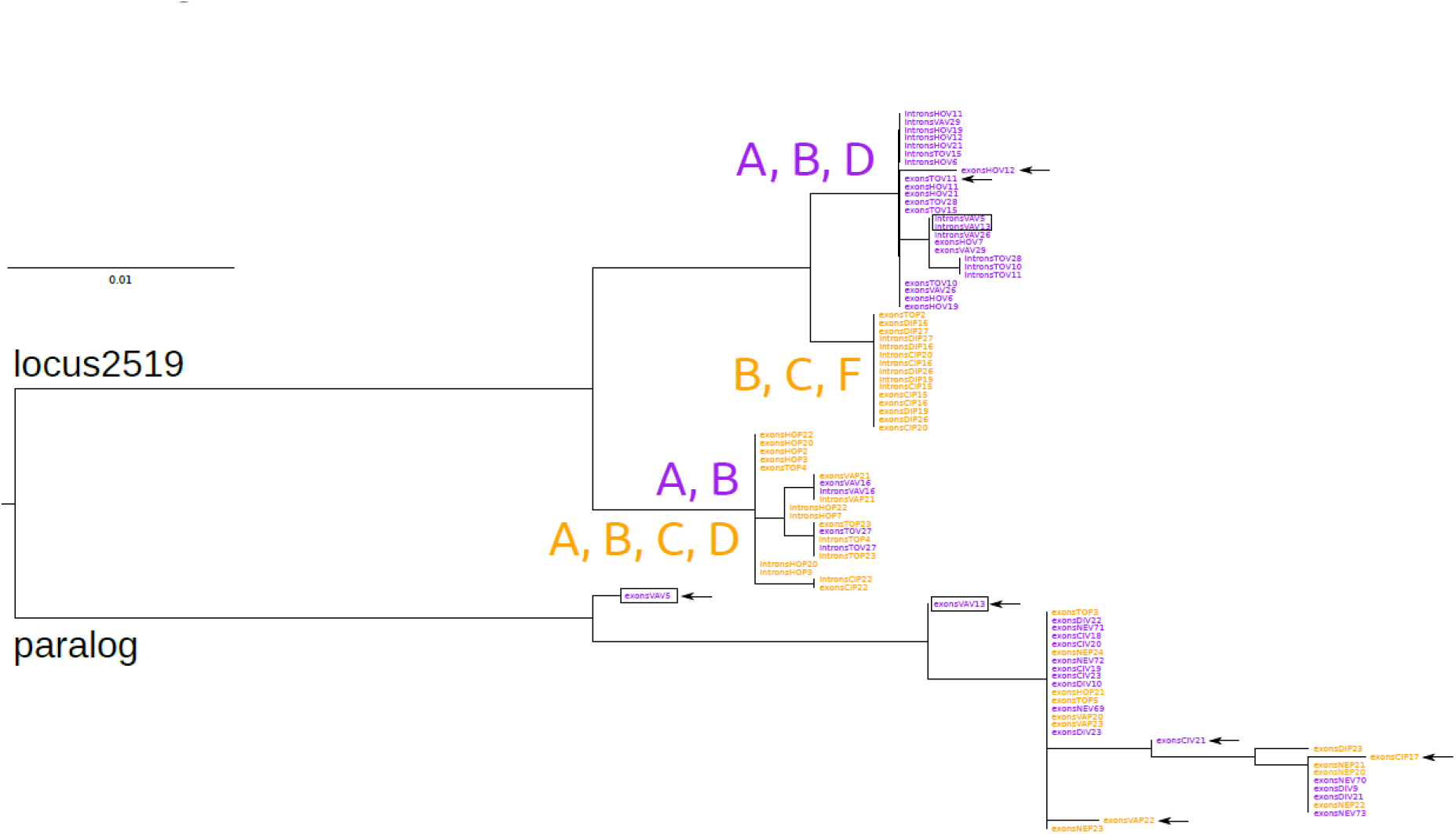
Maximum-likelihood tree of all sequences obtained from *Heliosperma pusillum* (orange) and *H. veselskyi* (purple) using different primer pairs of the putative outlier locus *locus2519* identified by Bayescan and its paralog. The labels "exons" or "introns" before the ID of each individual refers to the position of the primers used in the amplification (exons: both primers are in the coding region; introns: at least one of the two primers is in the non-coding regions). Sequences of the individuals marked with an arrow could not be properly interpreted because of the confusing signal of double amplification (always when using exons-based primers). Both paralogs have been sequenced for the individuals in black boxes. Population labels and species colours are as in Figure 1 in the main text. Individual labels correspond to Table S1.

## 8. Analysis of leaf microbiome

Besides the difference in the phyllosphere community, our metagenomic analysis highlighted also a difference in the private loci of either species assigned to flowering plants (Fig. S12, Tab. S2) and mapping to plastid sequences. Assuming a neutral accumulation of substitutions at the RADseq restriction enzyme cut-site, we expected a nearly equal proportion of allele drop-out in the two species (Gautier *et al*. 2013) and, hence, a similar proportion of private loci in the two species. The discrepancy in our results may be explained by a higher plastid genome proportion in the DNA extracted from *H. pusillum* individuals. An alternative explanation is related with the highly dynamic and massive mitochondrial genome found in several species of the closely related genus *Silene* (Sloan *et al*. 2014, Wu *et al*. 2015a). This mitochondrial genome has been described as fragmented in multiple circular chromosomes that can include several fast-evolving inserted copies of nuclear and plastid genes (Sloan & Wu 2014), and long intergenic regions that are actively transcribed and may have a regulatory activity (Wu *et al*. 2015b). More dedicated investigations are needed to confirm any difference in mitochondrial genome dynamics and structure between *H. pusillum* and *H. veselskyi*.

We also found a significant component of loci assigned to Diptera (in particular *Drosophila)* in four populations of *H. pusillum* (Figs. S10-S12, Tab S2). As *Drosophila* clearly prefers humid environments, it is not surprising it was only found on *H. pusillum* and not on the dry-adapted *H. veselskyi*. Why there is much less *Drosophila* contamination in E and F is less clear (all populations were sampled during the same summer season in July to August 2011). Further investigations are needed to identify the *Drosophila* species and clarify this aspect.

---

**Figure S9.**
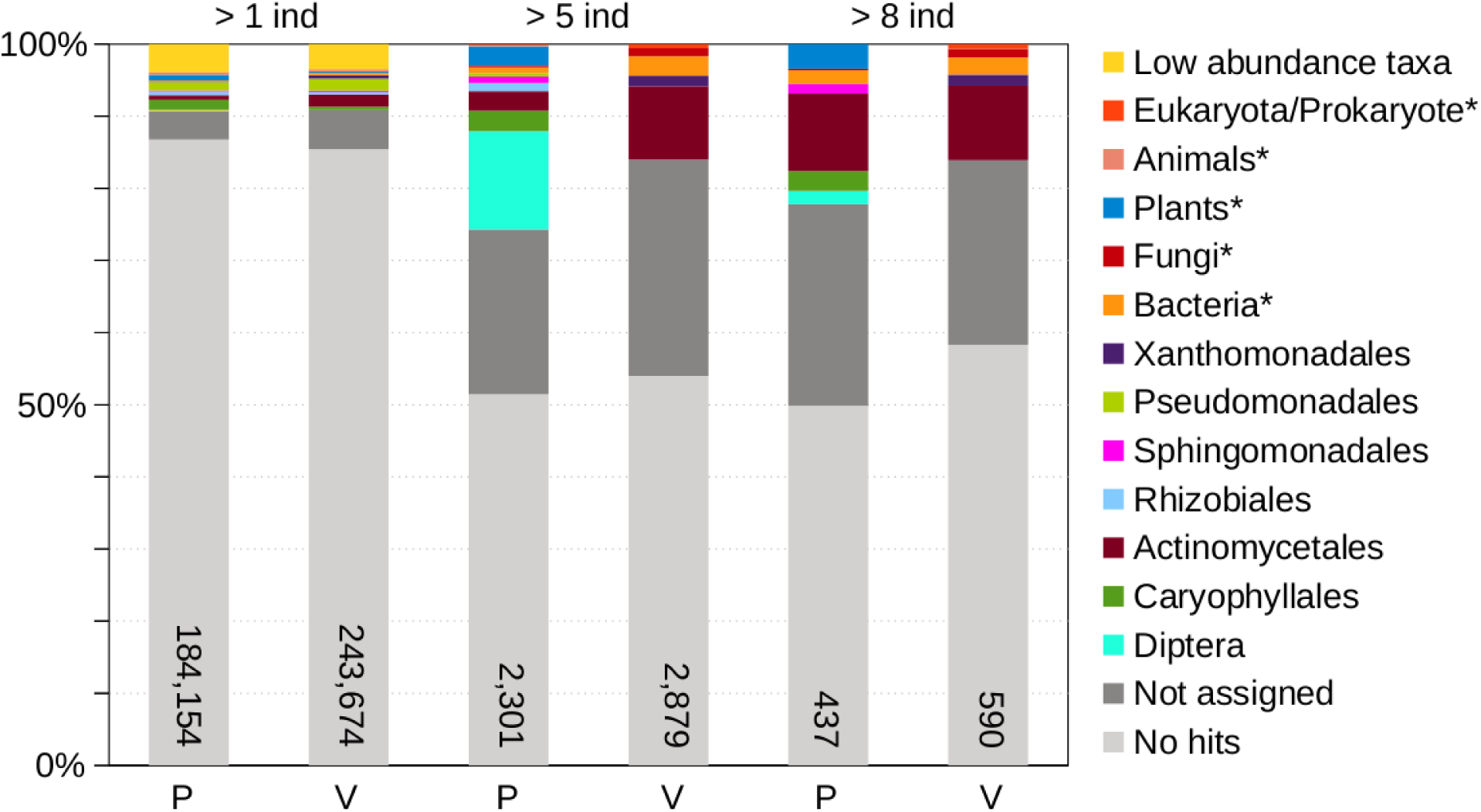
Proportion of loci blasting to the *nt* NCBI database and assigned to order level taxa in MEGAN (see the main text for details and reference) in three nested sets of loci selected as present in at least 1, 5, 8 individuals, respectively, of either *H. pusillum* or *H. veselskyi* (private loci of each species). Loci assigned to higher than order-like level taxa are also reported (asterisk). The number of loci included in each set are shown.

**Figure S10.**
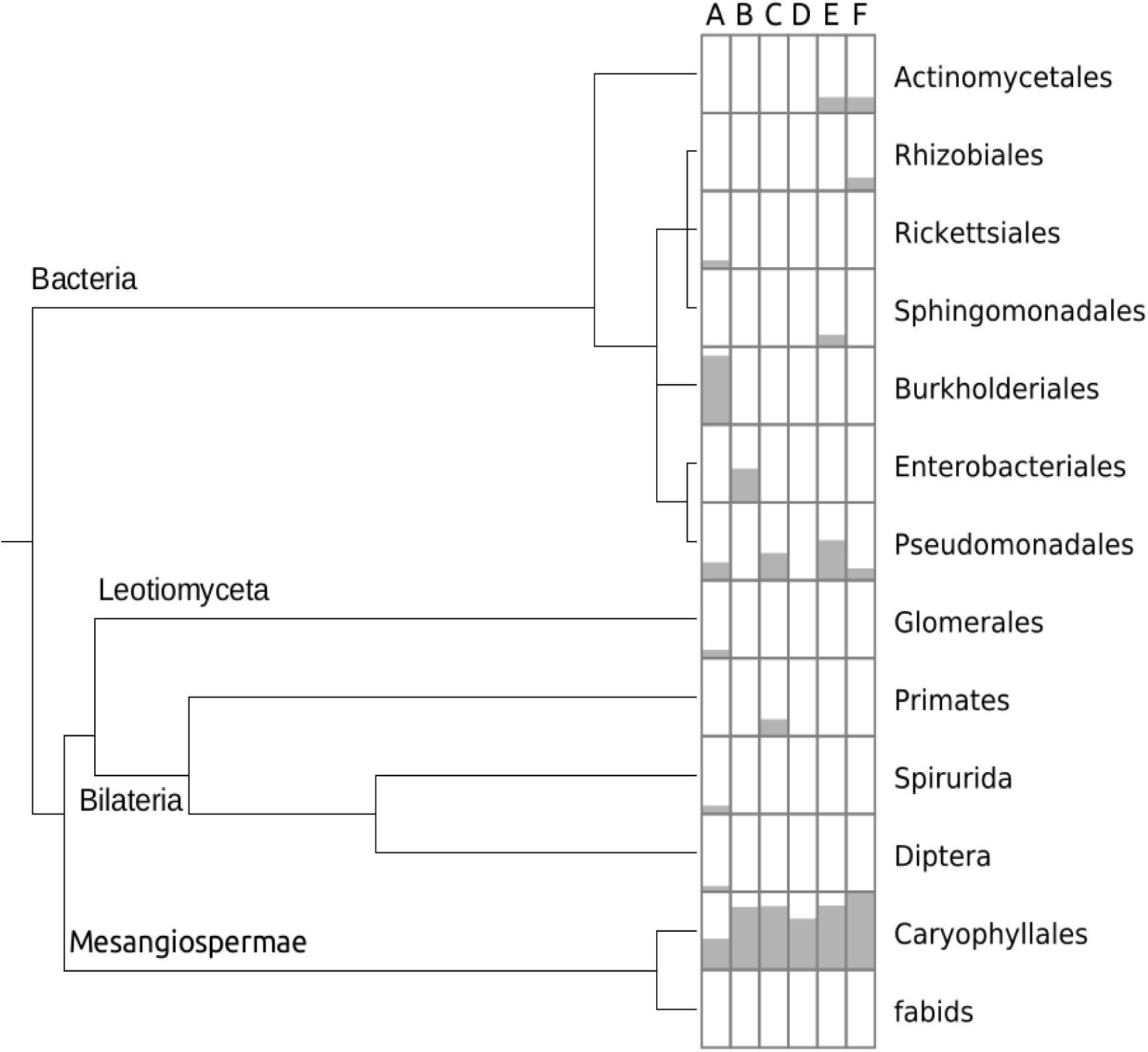
Shared loci between *Heliosperma pusillum* and *H. veselskyi* in each population pair. Loci were selected and retained for blasting if they are present in at least three individuals across the two species in each population pair. Most of the loci were expected to come from the target *Heliosperma* genome and not to be contaminants. Indeed, Caryophyllales (and all the higher hierarchy taxa up to Eukarya not shown here) are well represented by assigned hits. Very little of the exogenous DNA contamination is shared between the two species in each population pair. Taxa with the highest number of hits were *Variovorax paradoxus* in A, the only species contributing to the high prevalence of Burkholderiales in this locality, *Pseudomonas syringae* in A, *P. fluorescens* and *P. trivialis* in E, and *Rahnella aquatilis*, a quite rare Enterobacteriales, in B. Phylogenetic tree and nomenclature as automatically downloaded from GenBank NCBI database by MEGAN.

**Figure S11.**
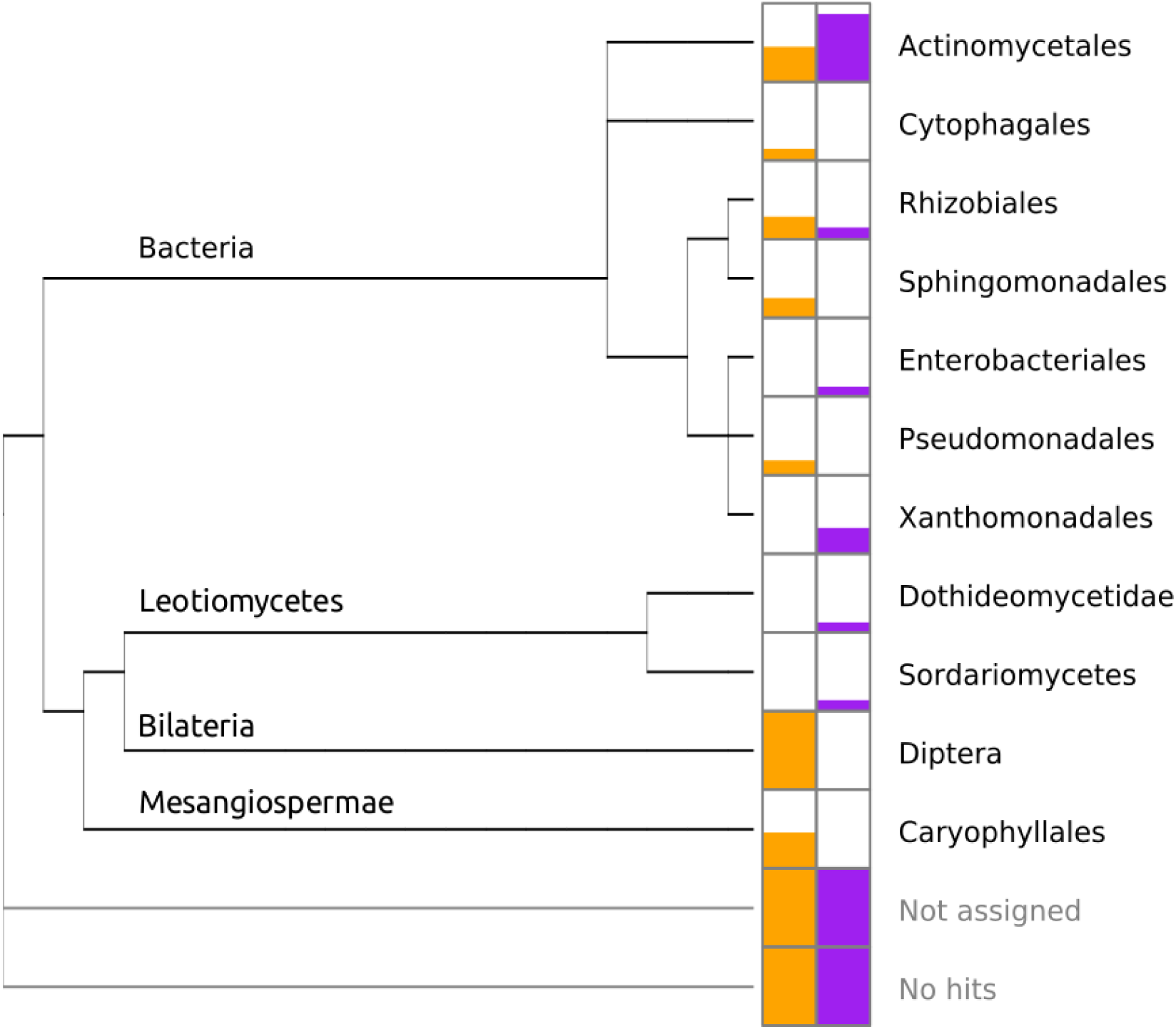
Normalized proportion of loci retrieved from populations of *Heliosperma pusillum* (orange) and *H. veselskyi* (purple) blasting to the *nt* NCBI database and assigned to order level taxa arranged on a phylogenetic tree (automatically downloaded from the NCBI database) by MEGAN (see main text for details and reference). A pruned version of this tree showing only the results for Bacteria and Leotiomycetes is in Figure 6 in the main text. Loci without hits to *nt* NCBI database or not assigned to a taxon by MEGAN are also shown.

**Table S3.**
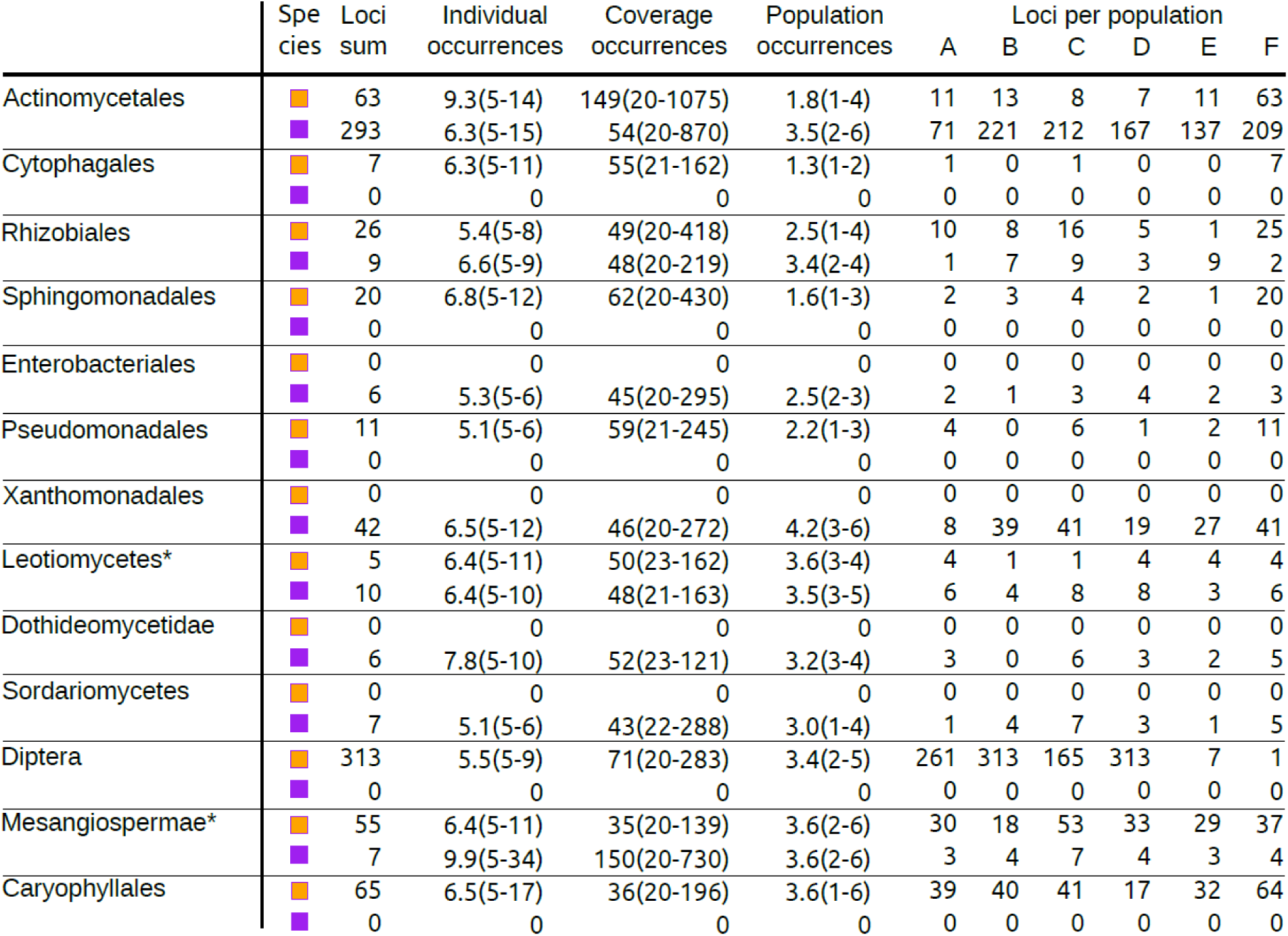
Number, occurrence across individuals, average coverage per individual, occurrence across populations (mean, min and max in brackets) of loci retrieved from populations of *Heliosperma pusillum* (orange) and *H. veselskyi* (purple) assigned to order level taxa as well as to higher-than-order-level taxa (marked by an asterisk) by MEGAN after blasting to the *nt* NCBI database (see Figure S11 and the main text for details). Number of loci present in each population are also shown. Population labels are as in Figure 1 in the main text. From 120 private *H. pusillum* loci assigned to Mesangiospermae and Caryophyllales, 108 were identified as plastid sequences, four as mitochondrial genes, three as cDNA clones, two as Copia Retrotransposon, one as ITS1, one as phospholipid transporting ATP-ase, and one as RNA-directed DNA polymerase homolog. From seven private *H. veselskyi* loci assigned to Mesangiospermae, four were identified as cDNA clones, one as auxin-response factor-like, one as microsatellite sequence, and one as 18S rRNA.

